# Plasma biomarkers of Alzheimer’s disease reveal conserved features of age-related neurodegeneration in chimpanzees and humans

**DOI:** 10.64898/2026.11.14.766572

**Authors:** Melissa K. Edler, Isha Mhatre-Winters, Chet C. Sherwood, Patrick R. Hof, Mary Ann Raghanti, Jason R. Richardson

## Abstract

Aging is a major risk factor for Alzheimer’s disease and related dementias (AD/ADRD). Yet, the physiological relationships linking age-related changes in peripheral biomarkers and measures of brain structure have only recently begun to be explored. Nonhuman primates, whose brains naturally express some of the pathologic hallmarks of AD/ADRD, including amyloid-beta (Aβ) plaques, vascular amyloid, and neurofibrillary tangles, represent a critical translational bridge between rodent models and humans. Here, we utilized ultrasensitive multiplex assays and structural MRI to examine the relationship of plasma AD/ADRD biomarkers with age, sex, and whole brain cortical thickness (CT) and gray matter volume (GMV) in a cohort of chimpanzees (*Pan troglodytes*), humans’ closest extant living relatives. Plasma levels (pg/ml) of total tau, phosphorylated tau 217 (pTau217), Aβ40, Aβ42, glial fibrillary acidic protein (GFAP), and neurofilament light chain (NfL) were measured in a cross-sectional sample of 154 chimpanzees ranging from 12-57 years of age. As in humans, aging was associated with increased plasma total tau, pTau217, Aβ40, Aβ42, GFAP, and NfL levels, as well as reduced whole brain CT and GMV in chimpanzees. The ratio of plasma Aβ42/Aβ40 increased with age in male chimpanzees, whereas aging was not significantly associated with the ratio of pTau217/Aβ42. Mediation analyses revealed that age-related increases in plasma Aβ42 mediated whole brain GMV atrophy in chimpanzees. Together, these findings demonstrate that plasma AD/ADRD biomarkers and measures of brain structure undergo age-related changes in chimpanzees, which is highly relevant to the study of human aging and AD/ADRD.

## Introduction

As the number of older people in the world grows quickly, the incidence rate and burden of Alzheimer’s disease and related dementias (AD/ADRD) increase at a rapid pace (World Health Organization, 2023). AD is the most common cause of dementia and is defined by irreversible changes in the brain that coincide with altered behavior and cognitive function. Primary neuropathologic hallmarks of AD are accumulation of amyloid beta (Aβ) protein in extracellular plaques and hyperphosphorylated tau protein in intraneuronal neurofibrillary tangles (NFT), which are accompanied by severe synaptic loss, dendritic atrophy, axonal deterioration, and widespread neuronal death in humans (Alzheimer, 1911; Arendt, 2009; Bussière, Giannakopoulos, et al., 2003; Bussière, Gold, et al., 2003; Cochran et al., 2014; West et al., 2004). In addition, more than 80% of human AD cases present with cerebral amyloid angiopathy (CAA), a condition in which Aβ deposits in and around the brain’s blood vessels (Arvanitakis et al., 2011; Jellinger, 2002). Presently, AD is confirmed by postmortem neuropathologic analyses, yet recent advances in fluid biomarker assays and *in vivo* neuroimaging are facilitating the identification of living individuals at risk for developing AD/ADRD, allowing for earlier diagnosis and intervention (Cohen et al., 2019; Grothe et al., 2021; Molinuevo et al., 2018).

A key obstacle in studying the preclinical prodromal phase of AD/ADRD is capturing subtle changes in imaging, fluid biomarkers, and brain function that occur over the course of several decades. The most often assessed AD/ADRD-associated fluid biomarkers include Aβ40, Aβ42, and phosphorylated tau (pTau) 181, 217, and 231 (Grothe et al., 2021; Salvadó et al., 2023). Other common biomarkers are neurofilament light protein (NfL), a measure of axonal injury and a proxy for neurodegeneration, and glial fibrillary acidic protein (GFAP), an indicator of astrogliosis and neuroinflammation (Ingannato et al., 2024; Schindler & Bateman, 2021; Simrén et al., 2021; Wang et al., 2024). During healthy aging in humans, plasma levels of Aβ40, Aβ42, total tau, pTau, GFAP, and NfL tend to increase, while the ratio of pTau217/Aβ42 remains stable and no change or a mild decrease with aging is observed in the ratio of Aβ42/Aβ40 (Arranz et al., 2024; Benussi et al., 2025; Cooper et al., 2024; Dage et al., 2016; De Meyer et al., 2024; González-Escalante et al., 2025; Lee et al., 2024; Lehmann et al., 2025; Lue et al., 2019; Mielke et al., 2021; T. Nakamura et al., 2018; Sun et al., 2025). In comparison, those with AD have higher levels of pTau, GFAP, NfL, and ratio of pTau217/Aβ42 but a lower ratio of Aβ42/Aβ40 than cognitively unimpaired subjects (Benedet et al., 2021; Mattsson et al., 2017; Mendes et al., 2024; Milà-Alomà et al., 2022; Santos et al., 2025; Schindler et al., 2019).

While rodents are the primary animal model in aging and AD/ADRD, they are unable to completely reproduce fluid biomarkers, neuroimaging, and pathologic profiles similar to humans (Drummond & Wisniewski, 2017; Dyrks et al., 1993; Goedert et al., 1989; Granzotto et al., 2024; Hodge et al., 2019; Quinn, 2018; Takuma et al., 2003). Nonhuman primates (NHP), such as chimpanzees, have more similar cognitive, communicative, and neuroanatomical features with humans compared to other commonly used models in aging and AD/ADRD research (Anderson & Gallup, 2011; Custance et al., 1995; de Waal, 1982; de Waal, 1996; Gardner & Gardner, 1969; Leavens & Racine, 2009; Leavens et al., 2009; Liebal et al., 2005; Lyn, 2012; Rumbaugh, 1977; Savage-Rumbaugh, 1986; Savage-Rumbaugh & Lewin, 1994; Schwartz & Beran, 2022; Sherwood et al., 2012; Tomasello & Call, 1997; Whiten et al., 1996). Moreover, chimpanzees show cross-sectional and longitudinal age-related changes in cognition and motor skill, and a substantial number of elderly chimpanzee brains spontaneously develop Aβ plaques, NFT, and severe CAA (Edler et al., 2023; Edler et al., 2020; Edler et al., 2018; Hopkins et al., 2021; Hopkins et al., 2019; Hopkins et al., 2009; Lacreuse et al., 2018; Lacreuse et al., 2014; Munger et al., 2019). Although geriatric chimpanzees and other great apes do not experience reductions in whole brain volume and weight like aged humans, region-specific reductions in cerebral gray matter volume (GMV) and cortical thickness (CT) have been noted in aged chimpanzees (Autrey et al., 2014; Hopkins et al., 2023, 2025; Hrvoj-Mihic et al., 2026; M. M. Mulholland et al., 2021).

Despite the similarities between human and ape brains, plasma biomarkers of aging and neurodegeneration have not been fully characterized in chimpanzees across the lifespan. Here, we quantified plasma levels of total tau, pTau217, Aβ40, Aβ42, GFAP, and NfL in a cross-sectional sample of 154 chimpanzees for whom archival whole brain GMV and CT were collected. Altogether, this dataset provides a unique opportunity to assess the effects of aging on plasma AD/ADRD biomarkers and their associations with measures of brain structure in chimpanzees.

## Materials and Methods

### Subjects

Archival plasma samples were collected between 2010 and 2013 during routine physical examinations from 154 chimpanzees ranging from 12 to 57 years old, housed at the Michale E. Keeling Center for Comparative Medicine and Research (KCCMR), which is part of The University of Texas MD Anderson Cancer Center (UTMDACC). Plasma samples were obtained from 64 males (mean age: 33.9 years, range: 14.9 – 51.8 years) and 90 females (mean age: 34.8 years, range: 12.2 – 57.3 years).

### Multiplex protein assays for plasma biomarkers

Blood samples were collected in EDTA tubes. Tubes were centrifuged at 1,300 x g for 10 minutes at room temperature (RT) and plasma was collected, aliquoted, and frozen at -80°C. Samples were thawed and centrifuged at 10,000 x g for 5 minutes at 4°C, and supernatants were aliquoted for each subsequent assay. Plasma biomarkers (pg/ml) were measured on the Meso Scale Discovery platform (MSD, Rockville, MD). Aβ40 and Aβ42 were quantified with the multiplex V-PLEX Aβ 6E10 panel 1 (K15200E-2). Ultrasensitive S-PLEX kits were used to measure total tau (K151APSS) and pTau217 (K151APFS), while GFAP and NfL levels were collected with custom S-PLEX human neurology panel 1 TURBO-BOOST kits (K151ANV-2). Vendor protocols were followed for each assay, and every sample was measured in duplicate. Assay analytics and limits of detection are provided in **Table S1**. Dilution linearity and spike recovery were assessed to confirm minimal matrix effects and an average recovery of analytes between 80 and 120 % (**Table S2**). Samples were run at a 1:4 dilution for the 6E10 (Aβ) assay, a neat dilution for total tau and pTau217, and a 1:2 dilution for GFAP and NfL. Plates were read on the MESO QuickPlex SQ 120 instrument and analyzed using the Discovery Workbench software. Concentrations were measured using an 8-point calibration curve with a 1/y^2^ weighted four-parameter logistic fit algorithm. A pooled plasma sample was added to each plate to assess intra-assay variation.

### Magnetic resonance imaging for cerebral gray matter volume (GMV) and cortical thickness (CT)

During the same timeframe that plasma samples were collected (2010-2013), magnetic resonance image (MRI) scans were obtained for chimpanzees housed at UTMDACC. Published measures of whole brain GMV (cc), including cortex, thalamus, basal ganglia, and cerebellum, and average whole brain CT (mm) from T1-structural MRIs were available for 154 individuals, including 64 males and 90 females (Hopkins et al., 2023; Vickery et al., 2020). The imaging sequence and post-image analyses for these scans have been described previously (Autrey et al., 2014; Hopkins et al., 2023; Vickery et al., 2020).

### Statistical analysis

Plasma biomarkers and ratios were converted from raw values (pg/ml) to natural logarithms (ln) due to a non-normal distribution, and outliers as confirmed by both ROUT and above the 99th percentile were removed prior to analyses (N=2/biomarker). Multiple linear regression modeling was conducted for age, sex, and interaction of age*sex effects on plasma biomarkers/ratios and measures of brain structure. When age was significantly associated with both plasma biomarkers/ratios and measures of brain structure, simple mediation analysis was employed using ordinary least squares regression and the Hayes Process macro (v.5.0) with bootstrapping (Model 4: 5,000) to examine if age alone or age-related changes in plasma levels affected measures of brain structure (**Fig. S1A**) (A. Hayes, 2018; A. F. Hayes, 2015). Moderated mediation (Model 7) was performed when a significant sex or age*sex interaction occurred in plasma biomarkers/ratios along with an age effect in measures of brain structure (i.e., CT and GMV), and variables were mean-centered (**Fig. S1B**). SPSS (v 31.0.0.0) and GraphPad PRISM (v 10.6.0) were utilized for all analyses and graphs. Significance was set at α ≤ 0.05.

## Results

Average plasma biomarker levels (pg/ml) are reported across the adult lifespan of chimpanzees (**Table S3**). Using raw values (pg/ml), ratios were also calculated for Aβ42/Aβ40, total tau/Aβ42, and pTau217/Aβ42 (**Table S4**), and mean CT (mm) and GMV (cc) are presented by sex and each decade of life (**Table S5**).

### Age and sex effects on plasma biomarkers, plasma ratios, and measures of brain structure

Multiple linear regression models with age, sex, and interaction of age*sex were utilized initially (**Table S6**), and the interaction term was removed from each model when non-significant (i.e., ≥ 0.05; **Table 1**). Age was positively correlated with all individual plasma biomarkers (**Fig. 1**) and the ratio of Aβ42/Aβ40 in chimpanzees (**Fig. 2**, **Table 1**). On the other hand, the ratio of total tau/Aβ42 significantly decreased with aging, pTau217/Aβ42 was not associated with age, and CT and GMV negatively correlated with advancing age (**Fig. 2-3**, **Table 1**).

**Table 1.**
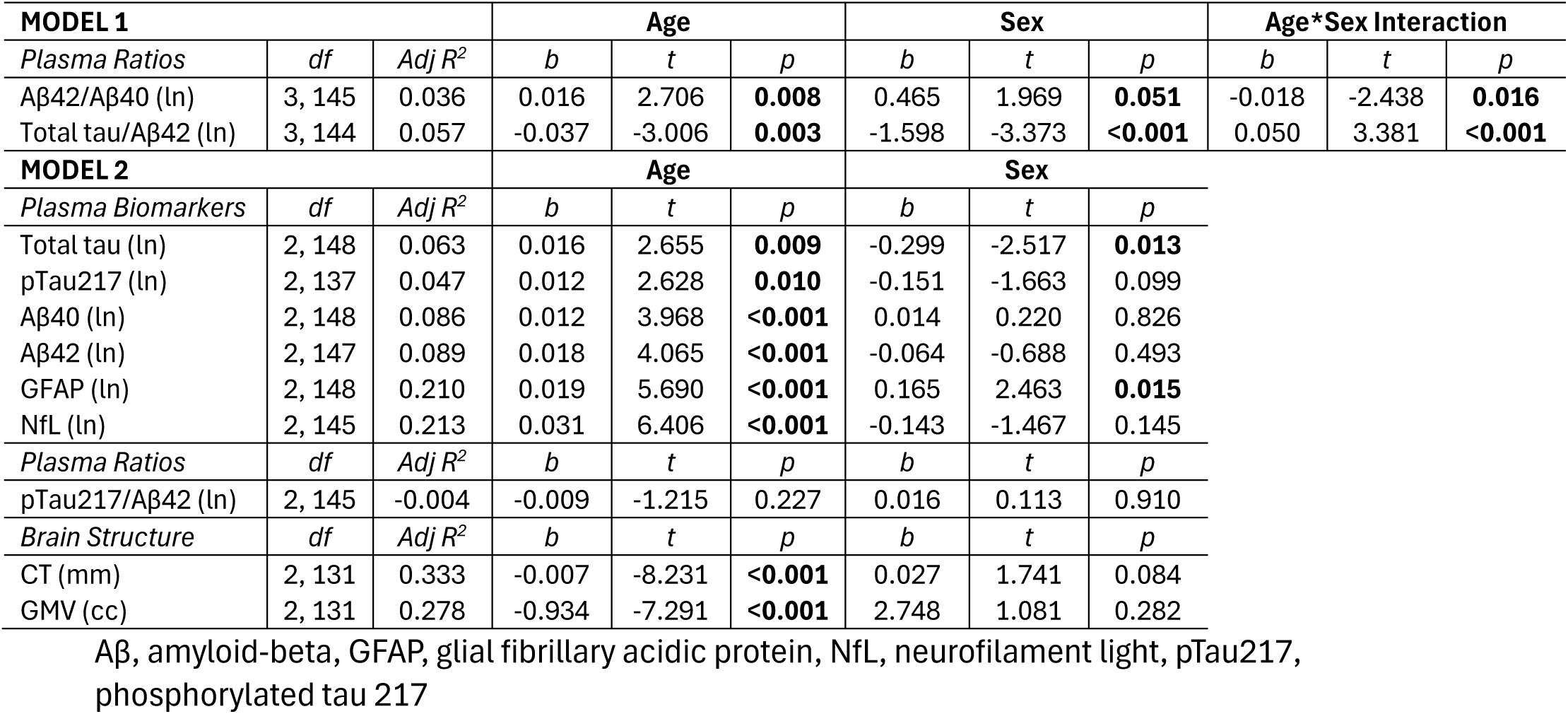
Final linear regression models for the main effects of age, sex, and interaction of age*sex with plasma biomarkers, plasma ratios, and measures of brain structure.

**Figure 1.**
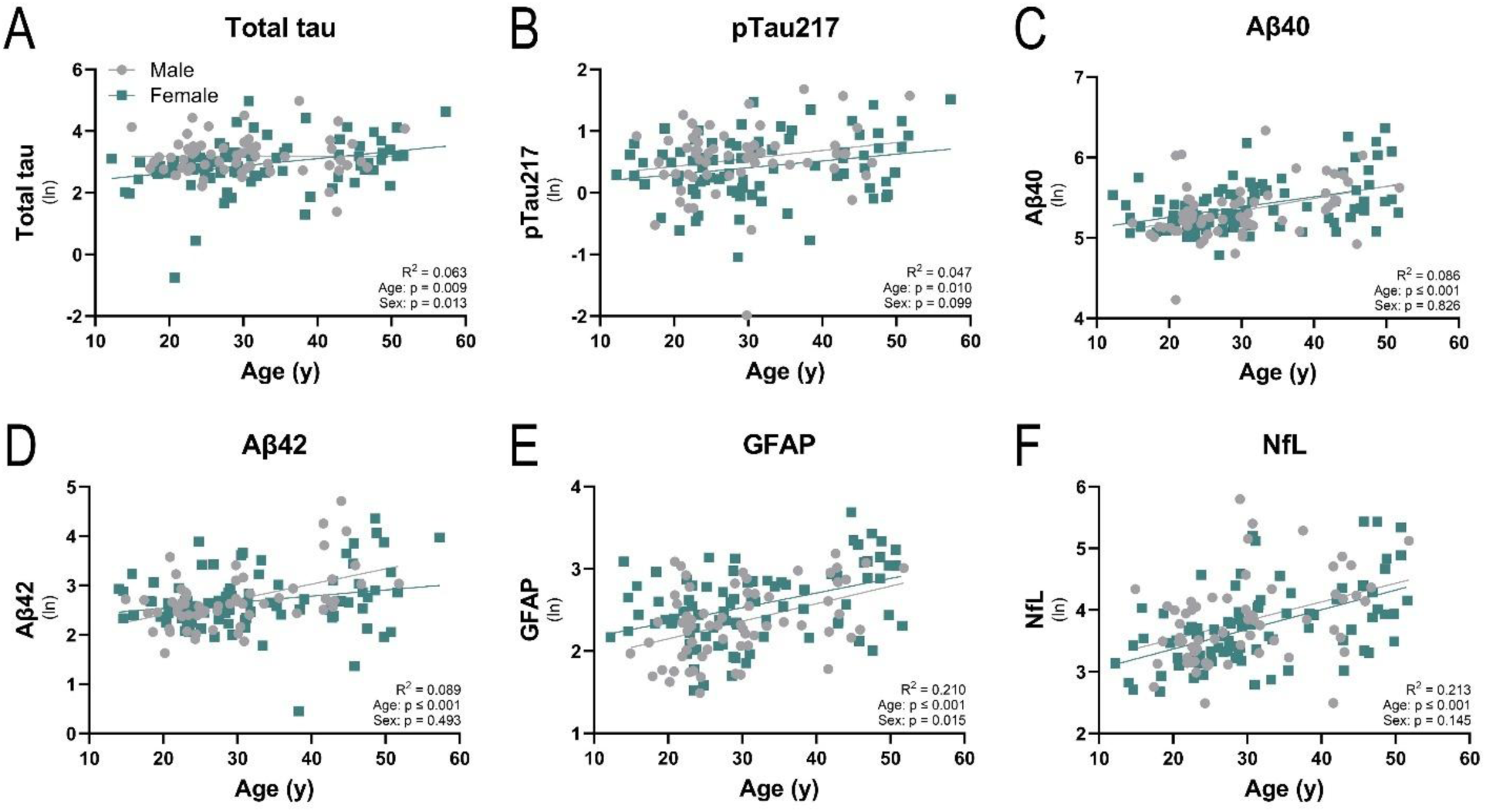
Association of age, sex (gray = male, pink = green), and plasma biomarkers (ln, natural logarithm) for total tau (A), phosphorylated tau 217 (pTau217; B), amyloid beta 40 (Aβ40; C) and 42 (Aβ42; D), glial fibrillary acidic protein (GFAP; E), and neurofilament light (NfL; F) in chimpanzees.

**Figure 2.**
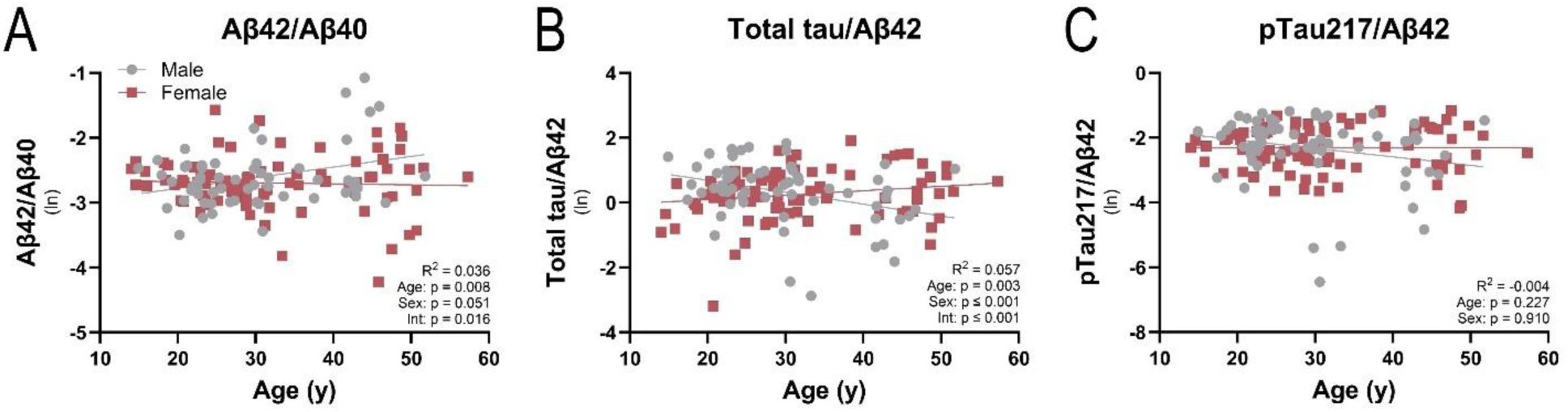
Association of age, sex (gray = male, red = female), and plasma ratios of Aβ42/Aβ40 (A), total tau/Aβ42 (B), and pTau217/Aβ42 (C) in chimpanzees.

**Figure 3.**
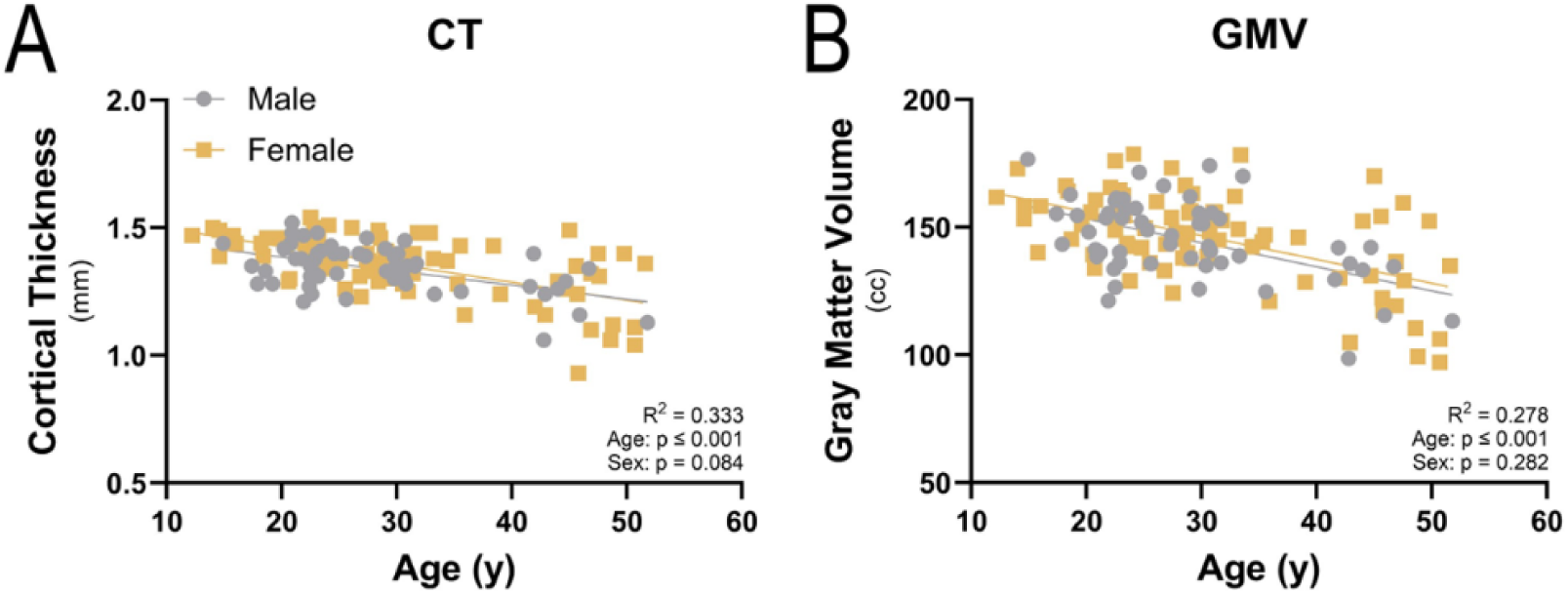
Association of age, sex (gray = male, yellow = female), cortical thickness (CT; A), and gray matter volume (GMV; B) in chimpanzees.

The main effect of sex was significant for total tau (**Fig. 1A**), GFAP (**Fig. 1E**), and ratios of Aβ42/Aβ40 (**Fig. 2A**) and total tau/Aβ42 (**Fig. 2B**, **Table 1**); moreover, two-way interactions between age*sex were significant for Aβ42/Aβ40 and total tau/Aβ42 (**Fig. 2A-B**). Females underwent a significant increase in plasma total tau with age (b = 0.023, p = 0.003), while levels remained stable in male chimpanzees across the lifespan (b = -0.0002, p = 0.981; **Fig. 1A**). Plasma GFAP was significantly higher in females compared to males, although age-related decreases were found in both sexes (male: b = 0.021, p = 0.004, female: b = 0.018, p ≤ 0.001; **Fig. 1E, Table S3**). The ratio of Aβ42/Aβ40 increased in male chimpanzees with age but not in females (male: b = 0.016, p = 0.010, female: b = -0.002, p = 0.698; **Fig. 2A**). Plasma total tau/Aβ42 significantly decreased with age in males only (male: b = -0.037, p = 0.007, female: b = 0.014, p = 0.093; **Fig. 2B**). No other sex differences or interactions between age*sex were noted in plasma biomarkers, ratios, or measures of brain structure (**Table 1**).

### Mediation analyses of age, plasma biomarkers/ratios, and measures of brain structure

To determine whether age alone or age-related changes in plasma biomarkers and ratios affected measures of brain structure independently, we employed simple mediation analyses. In the current analyses, age was treated as the independent or exposure variable (X), while measures of brain structure (i.e., CT and GMV) were the dependent or outcome variables (Y), and plasma biomarkers and ratios were evaluated as potential mediator variables (M) (**Fig. S1A**). Simple mediation (X→M→Y) was conducted when significant correlations were found between age and measures of brain structure (X→Y) and age and plasma biomarkers/ratios (X→M). Based on these conditions, mediator variables included pTau217, Aβ40, Aβ42, and NfL. Age alone was found to be a significant predictor with a negative direct effect on CT and GMV and a positive direct effect on pTau217, Aβ40, and NfL (**Table 2**). Additionally, an age-associated increase in plasma Aβ42 was a significant indirect mediator of GMV, explaining 14% of the overall effect as calculated by the ratio of indirect/total effect (p = 0.03; **Fig. 4**, **Table 3**). No other indirect effects on measures of brain structure were mediated by plasma biomarkers and ratios.

**Table 2.**
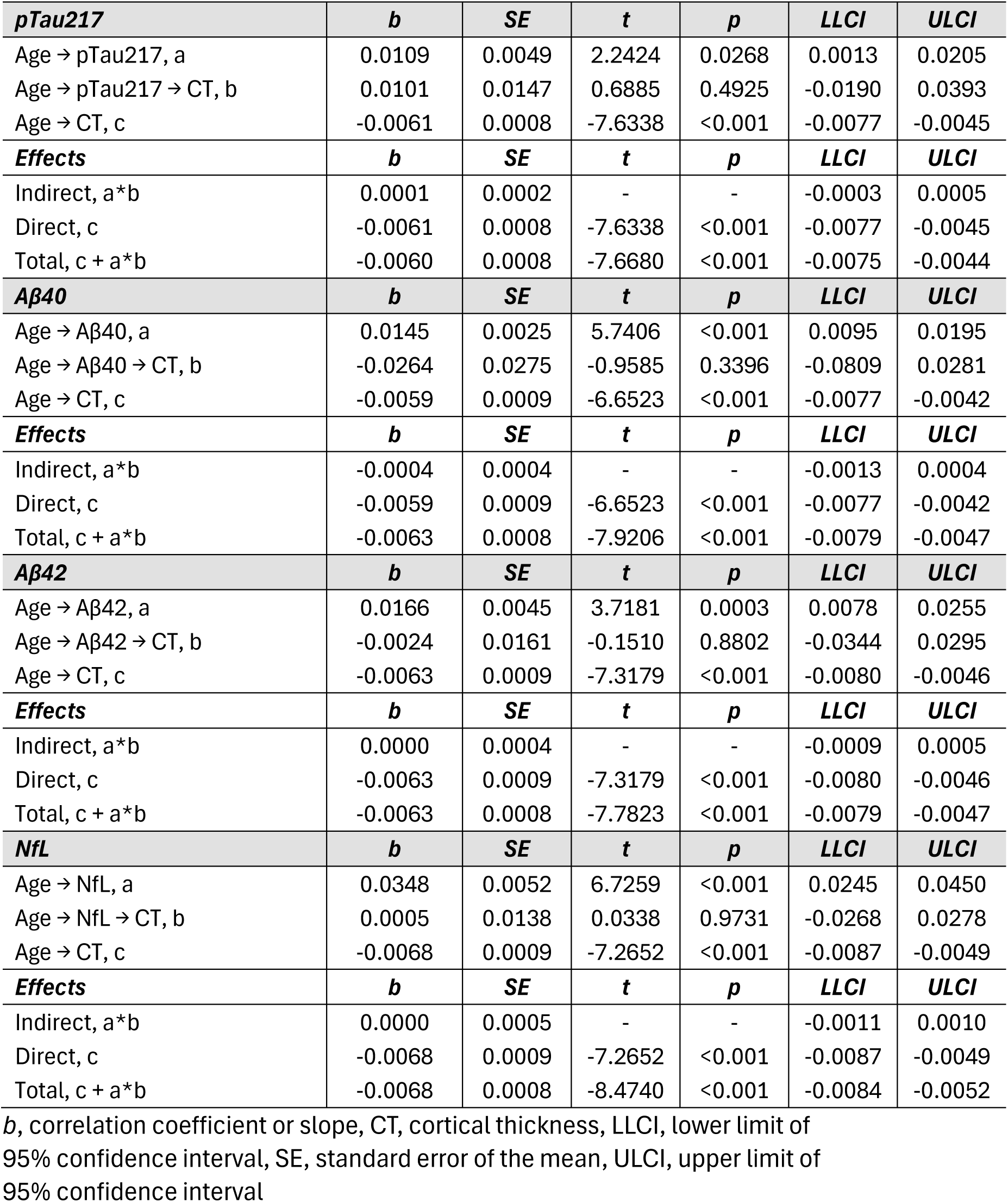
Simple mediation analysis of age (X) → plasma biomarkers/ratios (M) → CT (Y), where linear regression was used to measure the direct, indirect, and total effect of age on CT, as well as mediation by plasma biomarkers.

| ***pTau217*** | ***b*** | ***SE*** | ***t*** | ***p*** | ***LLCI*** | ***ULCI*** |
|---|---|---|---|---|---|---|
| Age → pTau217, a | 0.0109 | 0.0049 | 2.2424 | 0.0268 | 0.0013 | 0.0205 |
| Age → pTau217 → CT, b | 0.0101 | 0.0147 | 0.6885 | 0.4925 | -0.0190 | 0.0393 |
| Age → CT, c | -0.0061 | 0.0008 | -7.6338 | <0.001 | -0.0077 | -0.0045 |
| ***Effects*** | ***b*** | ***SE*** | ***t*** | ***p*** | ***LLCI*** | ***ULCI*** |
| Indirect, a*b | 0.0001 | 0.0002 | - | - | -0.0003 | 0.0005 |
| Direct, c | -0.0061 | 0.0008 | -7.6338 | <0.001 | -0.0077 | -0.0045 |
| Total, c + a*b | -0.0060 | 0.0008 | -7.6680 | <0.001 | -0.0075 | -0.0044 |
| ***Aβ40*** | ***b*** | ***SE*** | ***t*** | ***p*** | ***LLCI*** | ***ULCI*** |
| Age → Aβ40, a | 0.0145 | 0.0025 | 5.7406 | <0.001 | 0.0095 | 0.0195 |
| Age → Aβ40 → CT, b | -0.0264 | 0.0275 | -0.9585 | 0.3396 | -0.0809 | 0.0281 |
| Age → CT, c | -0.0059 | 0.0009 | -6.6523 | <0.001 | -0.0077 | -0.0042 |
| ***Effects*** | ***b*** | ***SE*** | ***t*** | ***p*** | ***LLCI*** | ***ULCI*** |
| Indirect, a*b | -0.0004 | 0.0004 | - | - | -0.0013 | 0.0004 |
| Direct, c | -0.0059 | 0.0009 | -6.6523 | <0.001 | -0.0077 | -0.0042 |
| Total, c + a*b | -0.0063 | 0.0008 | -7.9206 | <0.001 | -0.0079 | -0.0047 |
| ***Aβ42*** | ***b*** | ***SE*** | ***t*** | ***p*** | ***LLCI*** | ***ULCI*** |
| Age → Aβ42, a | 0.0166 | 0.0045 | 3.7181 | 0.0003 | 0.0078 | 0.0255 |
| Age → Aβ42 → CT, b | -0.0024 | 0.0161 | -0.1510 | 0.8802 | -0.0344 | 0.0295 |
| Age → CT, c | -0.0063 | 0.0009 | -7.3179 | <0.001 | -0.0080 | -0.0046 |
| ***Effects*** | ***b*** | ***SE*** | ***t*** | ***p*** | ***LLCI*** | ***ULCI*** |
| Indirect, a*b | 0.0000 | 0.0004 | - | - | -0.0009 | 0.0005 |
| Direct, c | -0.0063 | 0.0009 | -7.3179 | <0.001 | -0.0080 | -0.0046 |
| Total, c + a*b | -0.0063 | 0.0008 | -7.7823 | <0.001 | -0.0079 | -0.0047 |
| ***NfL*** | ***b*** | ***SE*** | ***t*** | ***p*** | ***LLCI*** | ***ULCI*** |
| Age → NfL, a | 0.0348 | 0.0052 | 6.7259 | <0.001 | 0.0245 | 0.0450 |
| Age → NfL → CT, b | 0.0005 | 0.0138 | 0.0338 | 0.9731 | -0.0268 | 0.0278 |
| Age → CT, c | -0.0068 | 0.0009 | -7.2652 | <0.001 | -0.0087 | -0.0049 |
| ***Effects*** | ***b*** | ***SE*** | ***t*** | ***p*** | ***LLCI*** | ***ULCI*** |
| Indirect, a*b | 0.0000 | 0.0005 | - | - | -0.0011 | 0.0010 |
| Direct, c | -0.0068 | 0.0009 | -7.2652 | <0.001 | -0.0087 | -0.0049 |
| Total, c + a*b | -0.0068 | 0.0008 | -8.4740 | <0.001 | -0.0084 | -0.0052 |

*b*, correlation coefficient or slope, CT, cortical thickness, LLCI, lower limit of 95% confidence interval, SE, standard error of the mean, ULCI, upper limit of 95% confidence interval

**Figure 4.**
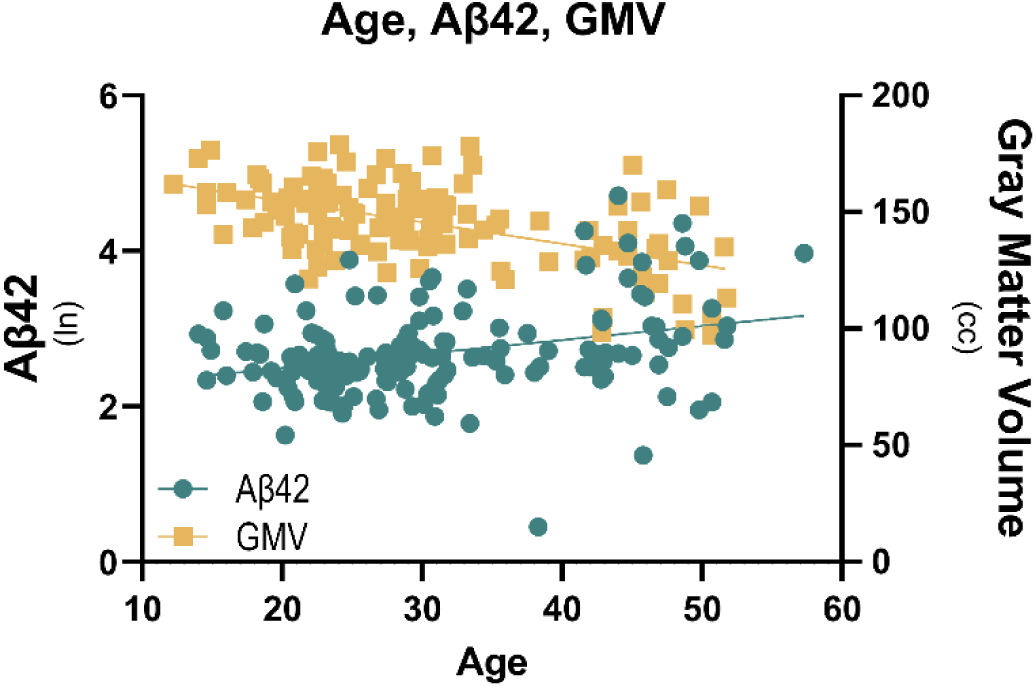
Mediational analyses revealed that age independently as well as age-related increases in plasma Aβ42 caused decreased gray matter volume in chimpanzees.

**Table 3.** Simple mediation analysis of age (X) → plasma biomarkers/ratios (M) → GMV (Y), where linear regression was used to measure the direct, indirect, and total effect of age on GMV, as well as mediation by plasma biomarkers.

| ***pTau217*** | ***b*** | ***SE*** | ***t*** | ***p*** | ***LLCI*** | ***ULCI*** |
|---|---|---|---|---|---|---|
| Age → pTau217, a | 0.0113 | 0.0048 | 2.3372 | 0.0211 | 0.0017 | 0.0208 |
| Age → pTau217 → GMV, b | 2.1846 | 2.5169 | 0.8680 | 0.3871 | -2.7987 | 7.1680 |
| Age → GMV, c | -0.9346 | 0.1365 | -6.8453 | <0.001 | -1.2049 | -0.6643 |
| ***Effects*** | ***b*** | ***SE*** | ***t*** | ***p*** | ***LLCI*** | ***ULCI*** |
| Indirect, a*b | 0.0246 | 0.0323 | - | - | -0.0442 | 0.0891 |
| Direct, c | -0.9346 | 0.1365 | -6.8453 | <0.001 | -1.2049 | -0.6643 |
| Total, c + a*b | -0.9100 | 0.1334 | -6.8206 | <0.001 | -1.1741 | -0.6458 |
| ***Aβ40*** | ***b*** | ***SE*** | ***t*** | ***p*** | ***LLCI*** | ***ULCI*** |
| Age → Aβ40, a | 0.0138 | 0.0024 | 5.6979 | <0.001 | 0.0090 | 0.0187 |
| Age → Aβ40 → GMV, b | -5.2610 | 4.6415 | -1.1335 | 0.2591 | -14.4443 | 3.9222 |
| Age → GMV, c | -0.8441 | 0.1437 | -5.8722 | <0.001 | -1.1285 | -0.5597 |
| ***Effects*** | ***b*** | ***SE*** | ***t*** | ***p*** | ***LLCI*** | ***ULCI*** |
| Indirect, a*b | -0.0728 | 0.0627 | - | - | -0.2052 | 0.0410 |
| Direct, c | -0.8441 | 0.1437 | -5.8722 | <0.001 | -1.1285 | -0.5597 |
| Total, c + a*b | -0.9169 | 0.1287 | -7.1233 | <0.001 | -1.1716 | -0.6623 |
| ***Aβ42*** | ***b*** | ***SE*** | ***t*** | ***p*** | ***LLCI*** | ***ULCI*** |
| Age → Aβ42, a | 0.0161 | 0.0045 | 3.5429 | 0.0006 | 0.0071 | 0.0251 |
| Age → Aβ42 → GMV, b | -5.6664 | 2.5095 | -2.2580 | **0.0257** | -10.6322 | -0.7006 |
| Age → GMV, c | -0.8290 | 0.1352 | -6.1333 | <0.001 | -1.0965 | -0.5616 |
| ***Effects*** | ***b*** | ***SE*** | ***t*** | ***p*** | ***LLCI*** | ***ULCI*** |
| Indirect, a*b | -0.0912 | 0.0585 | - | - | -0.2235 | -0.0017 |
| Direct, c | -0.8290 | 0.1352 | -6.1333 | <0.001 | -1.0965 | -0.5616 |
| Total, c + a*b | -0.9202 | 0.1310 | -7.0226 | <0.001 | -1.1795 | -0.6610 |
| ***NfL*** | ***b*** | ***SE*** | ***t*** | ***p*** | ***LLCI*** | ***ULCI*** |
| Age → NfL, a | 0.0350 | 0.0052 | 6.7628 | <0.001 | 0.0248 | 0.0453 |
| Age → NfL → GMV, b | -2.0225 | 2.1901 | -0.9235 | 0.3575 | -6.3567 | 2.3117 |
| Age → GMV, c | -0.9151 | 0.1492 | -6.1345 | <0.001 | -1.2103 | -0.6199 |
| ***Effects*** | ***b*** | ***SE*** | ***t*** | ***p*** | ***LLCI*** | ***ULCI*** |
| Indirect, a*b | -0.0709 | 0.0986 | - | - | -0.2947 | 0.1022 |
| Direct, c | -0.9151 | 0.1492 | -6.1345 | <0.001 | -1.2103 | -0.6199 |
| Total, c + a*b | -0.9860 | 0.1278 | -7.7130 | <0.001 | -1.2390 | -0.7330 |

*b*, correlation coefficient or slope, GMV, gray matter volume, LLCI, lower limit of 95% confidence interval, SE, standard error of the mean, ULCI, upper limit of 95% confidence interval

Moderated mediation was conducted when a significant sex effect or interaction of age*sex were observed in a mediator variable (i.e., plasma biomarkers/ratios). In our moderated mediation model, we tested for the indirect effect of age (X) on measures of brain structure (Y) via plasma biomarkers/ratios (M) with the indirect effect being moderated by sex (W, **Fig. S1B**). Mediator variables for these analyses included total tau, GFAP, and the ratios of Aβ42/Aβ40 and total tau/Aβ42. The conditional indirect effect of sex was non-significant for all investigated measures of brain structure and plasma biomarkers/ratios, and the index of moderated mediation, which quantifies the degree to which the indirect effect is moderated by sex, is reported for each mediator variable by measures of brain structure (**Table 4**).

**Table 4.** Index of moderated mediation and lower (LLCI) and upper limit of 95% confidence interval (ULCI) with age as the independent variable (X), measures of brain structure as the dependent variable (Y), plasma biomarkers/ratios as the mediator variables (M), and sex as the moderator variable (W). If the null (i.e., 0) falls between the lower and upper confidence intervals, the moderation effect of sex is non-significant.

| ***CT*** | ***Index*** | ***LLCI*** | ***ULCI*** |
|---|---|---|---|
| Total Tau | 0.0000 | -0.0005 | 0.0006 |
| GFAP | -0.0001 | -0.0006 | 0.0005 |
| Aβ42/Aβ40 | -0.0001 | -0.0016 | 0.0010 |
| Total Tau/Aβ42 | 0.0002 | -0.0007 | 0.0012 |
| ***GMV*** | ***Index*** | ***LLCI*** | ***ULCI*** |
| Total Tau | -0.0018 | -0.0775 | 0.1020 |
| GFAP | -0.0117 | -0.0979 | 0.0524 |
| Aβ42/Aβ40 | 0.1193 | -0.0333 | 0.3371 |
| Total Tau/Aβ42 | 0.0915 | -0.0300 | 0.2467 |

### Covariation between plasma biomarkers

Partial correlation coefficients between plasma biomarkers while controlling for age and sex are reported in **Fig. S2** and **Table S10**. All plasma biomarkers were significantly and positively correlated with each other except total tau with GFAP (p = 0.21) and pTau217 with Aβ42 (p = 0.69).

## Discussion

In this study, we investigated the relationship of age, sex, AD-related plasma biomarkers, and measures of brain structure in a large cohort of chimpanzees. Aging was associated with increased levels of all examined plasma biomarkers (**Fig. 1**), reduced CT (**Fig. 3A**), and whole brain gray matter atrophy (**Fig. 3B**, **Table 1**). Furthermore, mediation analysis revealed age-induced increases in plasma Aβ42 contributed to decreased overall GMV in chimpanzees (**Fig. 4**, **Table 3**). Sex differences were noted with females undergoing an age-related increase in total tau (**Fig. 1A**) as well as having higher GFAP (**Fig. 1E**), while males exhibited a rise in Aβ42/Aβ40 (**Fig. 2A**) and a reduction in total tau/Aβ42 with aging (**Fig. 2B**, **Table 1**). Below, we will review each measure in a comparative context.

### Sex differences in plasma biomarkers and ratios

Some sex differences were noted in plasma biomarkers and ratios in chimpanzees. Female chimpanzees displayed an age-related increase in total tau (**Fig. 1A**) and had higher GFAP levels than males (**Fig. 1E**). Our findings are comparable to reports in human females, which found greater plasma GFAP concentrations and a higher increase in plasma total tau concentrations over time compared to men (Baldacci et al., 2020; Labonte et al., 2025). Male chimpanzees exhibited a rise in Aβ42/Aβ40 (**Fig. 2A**) and a reduction in total tau/Aβ42 with aging (**Fig. 2B**, **Table 1**). Both male and female chimpanzees displayed age-related increases in plasma Aβ40 and Aβ42 (**Fig. 1**, **Table 1**). While the age-associated slope was similar between sexes for Aβ40 (male: B = 0.015, p = 0.001; female: B = 0.013, p ≤ 0.0001), it differed for Aβ42 with male chimpanzees exhibiting a two-fold higher slope (male: B = 0.031, p ≤ 0.0001; female: B = 0.012, p =0.035), which likely contributed to the age-related increase in the ratio of Aβ42/Aβ40 in males only. This result may indicate a decreased risk for developing AD lesions in male chimpanzees, as low plasma levels of Aβ42/Aβ40 are associated with increased risk of cortical Aβ burden and cognitive decline in humans (Fandos et al., 2017; Seppala et al., 2010). Indeed, the number of female chimpanzees in this study with amyloid or tau pathology far outweighed the number of male chimpanzees, although sex differences in longevity likely contributed as well as the known higher risk of developing AD in female humans.

### Aging and AD/ADRD-related plasma biomarkers

In the past decade, a growing number of studies have examined AD/ADRD-related plasma biomarkers in humans, but investigating these biomarkers across the lifespan of NHP remains understudied. Plasma total tau and pTau217 concentrations in humans generally rise with aging, and elevated pTau217 levels in cognitively unimpaired individuals are a strong indicator of future clinical progression to mild cognitive impairment (MCI), a prodromal stage of AD/ADRD (J. Chen et al., 2023; Dage et al., 2016; Lee et al., 2024; Lue et al., 2019; Sun et al., 2025). Like in humans and in a preliminary report of serum biomarkers in chimpanzees, we found age-related increases in plasma total tau and pTau217 in chimpanzees (**Fig. 1A-B**, **Table 1**) (M. Mulholland et al., 2026). In contrast, total tau plasma concentrations in rhesus macaques (*Macaca mulatta*, maximum age = 22 years), as measured by a MSD assay validated for use in NHP, did not exhibit increases during aging (Metzler et al., 2025). Parallel to chimpanzees and humans, the specific measure of plasma pTau217 by nanoneedle technology increased across the lifespan of rhesus macaques (Datta et al., 2024). Another recently published report using NULISA-seq supports this finding, with an age-related increase in plasma pTau217 between adult and geriatric rhesus macaques (maximum age = 34 years) (Diniz et al., 2026), although a preliminary study using the same platform found an absence of association with plasma pTau217 and aging in rhesus macaques (maximum age = 29 years) but an increase with advancing age in baboons (maximum age = 24 years) (M. M. Mulholland et al., 2026). Differences in cohort size, age range, and sex distribution may contribute to these findings. Additionally, assay performance in NHP samples also may play a role and warrants further validation beyond analyte detectability and antibody sequence homology.

We also identified age-related increases in plasma Aβ40 and Aβ42 in chimpanzees, as is the case for most studies in cognitively unimpaired and Aβ-negative humans and in a preliminary study of chimpanzee serum (**Fig. 1C-D**, **Table 1**) (Cecchetti et al., 2024; J. Chen et al., 2023; Lehmann et al., 2025; Lue et al., 2019; M. Mulholland et al., 2026; T. Nakamura et al., 2018; Sun et al., 2025; Zecca et al., 2021). Previous reports identified similar age-based increases in plasma Aβ40 levels in coppery titi monkeys (*Plecturocebus cupreus*) and gray mouse lemurs (*Microcebus murinus*), although these assays (MSD 4G8, Invitrogen Aβ40), although species-specific validation was not reported (Gary et al., 2018; Rothwell et al., 2024; Roy et al., 2015). Published and preliminary studies in rhesus macaques vary with an absence of age-related effects or an increase with age noted (Diniz et al., 2026; Metzler et al., 2025; M. M. Mulholland et al., 2026). Conversely, decreased plasma Aβ42 was observed between female rhesus macaque cohorts of ≤5-year and 5-7-year-olds, but this difference was not viewed among the 8-16 or 17-22-year-old groups, while other reports found no association with age (Metzler et al., 2025). The youngest ape in the current study was 12 years old, so it remains unclear if an analogous developmental age-based decline in plasma Aβ42 occurs in chimpanzees. Age-based increases were reported in plasma Aβ40 and Aβ42 in baboons in an unpublished investigation (M. M. Mulholland et al., 2026).

In addition, we reported greater plasma GFAP and NfL levels with advancing age in chimpanzees (**Fig. 1E-F**, **Table 1**), like accounts in humans, coppery titi monkeys, and common marmosets (*Callithrix jacchus*) (Cooper et al., 2024; De Meyer et al., 2024; González-Escalante et al., 2025; Rothwell et al., 2024; Sun et al., 2025). In rhesus macaques using MSD assays for NHP, plasma NfL increased during aging, particularly in middle adulthood, while plasma GFAP decreased from the pre-adult (1-5 years) to young adult (5-7 years) group in males only (Metzler et al., 2025). Other reports based on screening assays in rhesus macaques indicated age-related increases in both NfL and GFAP (Diniz et al., 2026; M. M. Mulholland et al., 2026). A pre-print reported that in baboons, aging was associated with increased plasma NfL, but not GFAP (M. M. Mulholland et al., 2026). Overall, this evidence indicates that future work is needed to identify the shared and unique fluid biomarker profiles for each NHP species, although chimpanzees appear to be the closest to modeling human aging with age-associated modifications in AD/ADRD-related plasma biomarker profiles in chimpanzees corresponding well to cognitively unimpaired elderly humans.

### Aging and AD/ADRD-related plasma biomarker ratios

In recent years, ratios of plasma biomarkers have shown promise over individual biomarkers in the diagnosis of AD/ADRD, yet much remains unknown about whether these ratios are altered during the lifespan of humans and NHP. The most frequently assessed plasma ratios include Aβ42/Aβ40 and pTau217/Aβ42, which are FDA-approved blood tests for individuals over the age of 55 years. Plasma Aβ42/Aβ40 correlates with cerebrospinal fluid (CSF) Aβ42/Aβ40 levels and Aβ-positive positron emission tomography (PET) imaging, and it can identify individuals with abnormal brain Aβ burden or those at high risk for future conversion to Aβ-PET positivity with relatively high accuracy (A. Nakamura et al., 2018; Schindler et al., 2019). Studies performed in cognitively unimpaired human subjects and Aβ-negative individuals, ranging from 45-89 years old, noted either a decreased plasma ratio of Aβ42/Aβ40 or no change with aging (Arranz et al., 2024; J. Chen et al., 2023; De Meyer et al., 2024; González-Escalante et al., 2025; Lehmann et al., 2025; Sun et al., 2025). In contrast, we observed an age-related increase in the Aβ42/Aβ40 ratio in male chimpanzees, while the ratio remained stable in females during aging (**Fig. 2A**, **Table 1**).

The variation in Aβ42/Aβ40 plasma ratio between species could be a crucial factor contributing to the significant NFT load observed in humans with AD compared to the relatively minimal tau lesions despite extensive Aβ pathology seen in chimpanzees and other NHP (Freire-Cobo et al., 2021; Heuer et al., 2012). Decreased Aβ42/Aβ40 plasma ratio is associated with greater cortical Aβ burden in cognitively unimpaired, MCI, and AD individuals (Fandos et al., 2017; Trelle et al., 2025). While aging alone is associated with increased amyloid lesions in the brain, PET-amyloid positivity, and neuroinflammation in humans, the effect of age on tau deposition relies on Aβ load, which leads to greater tau burden at a younger age and worse cognitive outcomes in late-onset AD (Ottoy et al., 2020; Zhang et al., 2023). In contrast, Aβ42/Aβ40 plasma ratio remained stable across the lifespan of female chimpanzees and increased with age in male chimpanzees, which could reflect age-induced, species-specific alterations in plasma Aβ40 and Aβ42 levels or different rates of cortical Aβ42 accumulation between species.

Although the plasma Aβ42/Aβ40 ratio reliably detects brain PET-amyloid positivity, pTau biomarkers correlate better with cognitive deficits and are more sensitive in determining those likely to progress to AD (Cousins et al., 2025; Dage et al., 2016). To date, plasma pTau217 and the pTau217/Aβ42 ratio have the highest sensitivity for detecting Aβ-PET positivity and brain amyloid levels (González-Escalante et al., 2025; Lehmann et al., 2025). Studies in humans have largely reported a lack of change in plasma pTau217/Aβ42 in cognitively unimpaired subjects during aging (age range: 50-85 years), and here, we report a similar stability in the plasma ratio of pTau217/Aβ42 in our cohort of chimpanzees across the lifespan (Arranz et al., 2024; Cecchetti et al., 2024; Lehmann et al., 2025).

### Aging and measures of brain structure

Structural MRI studies in humans have identified global age-related gray matter loss, although some research noted an absence of volumetric atrophy in cognitively healthy older adults (Abe et al., 2008; Allen et al., 2005; Burgmans et al., 2009; Lemaitre et al., 2012; Neufeld et al., 2022; Pfefferbaum et al., 1994; Ramanoël et al., 2018; Sowell et al., 2003). Cortical thinning also has been observed across the human lifespan, with models demonstrating a steep decline in CT after age 60 (Lemaitre et al., 2012; Potvin et al., 2017; Raz et al., 1997; Salat et al., 2004; Storsve et al., 2014). In the current investigation, we observed analogous age-related losses in GMV and CT as have been previously reported in chimpanzees (X. Chen et al., 2013; Hopkins et al., 2023, 2025; Lacreuse et al., 2020; M. M. Mulholland et al., 2021; Vickery et al., 2020).

### AD/ADRD-related plasma biomarkers, ratios, and measures of brain structure

In this study, we explored whether age-associated changes in AD/ADRD-related plasma biomarkers were associated with GMV and CT. Importantly, plasma biomarkers were not found to mediate the effects of age on CT in chimpanzees (**Table 2**), which is consistent with outcomes in humans where advanced age, but not plasma concentrations of total tau, pTau217, GFAP, NfL, or Aβ42/Aβ40, were correlated with reduced CT (Asken et al., 2022; Brodman et al., 2025; Chiu et al., 2017). However, one report determined that plasma pTau217 was a significant predictor of temporal cortical thinning in non-demented humans (Pereira et al., 2021). Conversely, we observed that age-related increases in plasma Aβ42 promoted GMV atrophy in chimpanzees, which supports a recent preliminary report that found age-based associations with serum biomarkers and GMV in chimpanzees (**Fig. 4**, **Table 3**) (M. Mulholland et al., 2026). This discovery differs from reports in NHP and humans. In humans, a population-based study noted that lower plasma Aβ42 levels were associated with smaller hippocampal volume, but not with total gray or white matter volumes (Hilal et al., 2018). Likewise, a low baseline ratio of Aβ42/Aβ40 in individuals with subjective cognitive decline was accompanied by decreased hippocampal and cortical volumes (Fandos et al., 2017). Plasma pTau217 and NfL were found to be predictive of hippocampal volume loss in non-demented individuals, whereas we did not observe an association between plasma total tau or pTau217 levels and whole brain GMV in chimpanzees (Pereira et al., 2021; Saloner et al., 2024). Other reports found a negative correlation with GMV and plasma NfL in a sample of healthy adults (N=173: >60 years), while high plasma GFAP concentrations were related to lower total white matter volume, but not GMV, in normal older adults with mild cognitive changes (N=50: mean age: 72.7 y) (Asken et al., 2022; Karoly et al., 2021). Most analyses of associations between measures of brain structure and plasma biomarkers in humans focused on regional, rather than global, changes, and similar prospective investigations in NHP are required.

### AD/ADRD-related neuropathology with plasma biomarker ratios and measures of brain structure

Identifying fluid biomarkers that correlate with the burden of Aβ and tau deposition in the brain and predict future cognitive decline is imperative in the prevention of AD/ADRD and developing effective interventions for the prodromal phase of disease. Studies of AD/ADRD plasma biomarkers in NHP with confirmed postmortem AD neuropathology are incredibly rare. A report in older gray mouse lemurs observed high plasma Aβ40 levels in animals with and without amyloid plaques (Roy et al., 2015). Here, we provide raw data for plasma biomarkers (**Fig. 5**, **Table S7**), plasma ratios (**Fig.6, Table S8**), and measures of brain structure (**Fig. 7, Table SG**) in 22 chimpanzees within this cohort who died after 2013 from natural causes or were euthanized for quality-of-life reasons, and subsequent neuropathologic analyses were performed for Aβ (i.e., Aβ42-immunoreactive plaques and CAA) and tau (i.e., AT8-immunoreactive NFT) in formalin-fixed brains (Edler et al., 2023; Edler et al., 2017). These chimpanzees were categorized by the presence (+) or absence (-) of neocortical moderate/severe CAA in three or more layers (A) and NFT (T) into three cohorts including those with CAA only (A+T-, N=7: 2 males, mean age=36.7 years, 5 females, mean age=33.1 years), those with CAA and NFT (A+T+, N=4: 4 females, mean age=49.6 years), and those exhibiting minimal-to-mild neocortical CAA and Aβ plaques limited to one or two layers and an absence of neocortical NFT (A-T-, N=11: 6 males, mean age=31.2 years, 5 females, mean age=27.4 years). Statistical analyses were not performed due to small sample size. Contrary to what has been observed in humans, no obvious trends were noted between chimpanzees with or without AD lesions in their brains.

**Figure 5.**
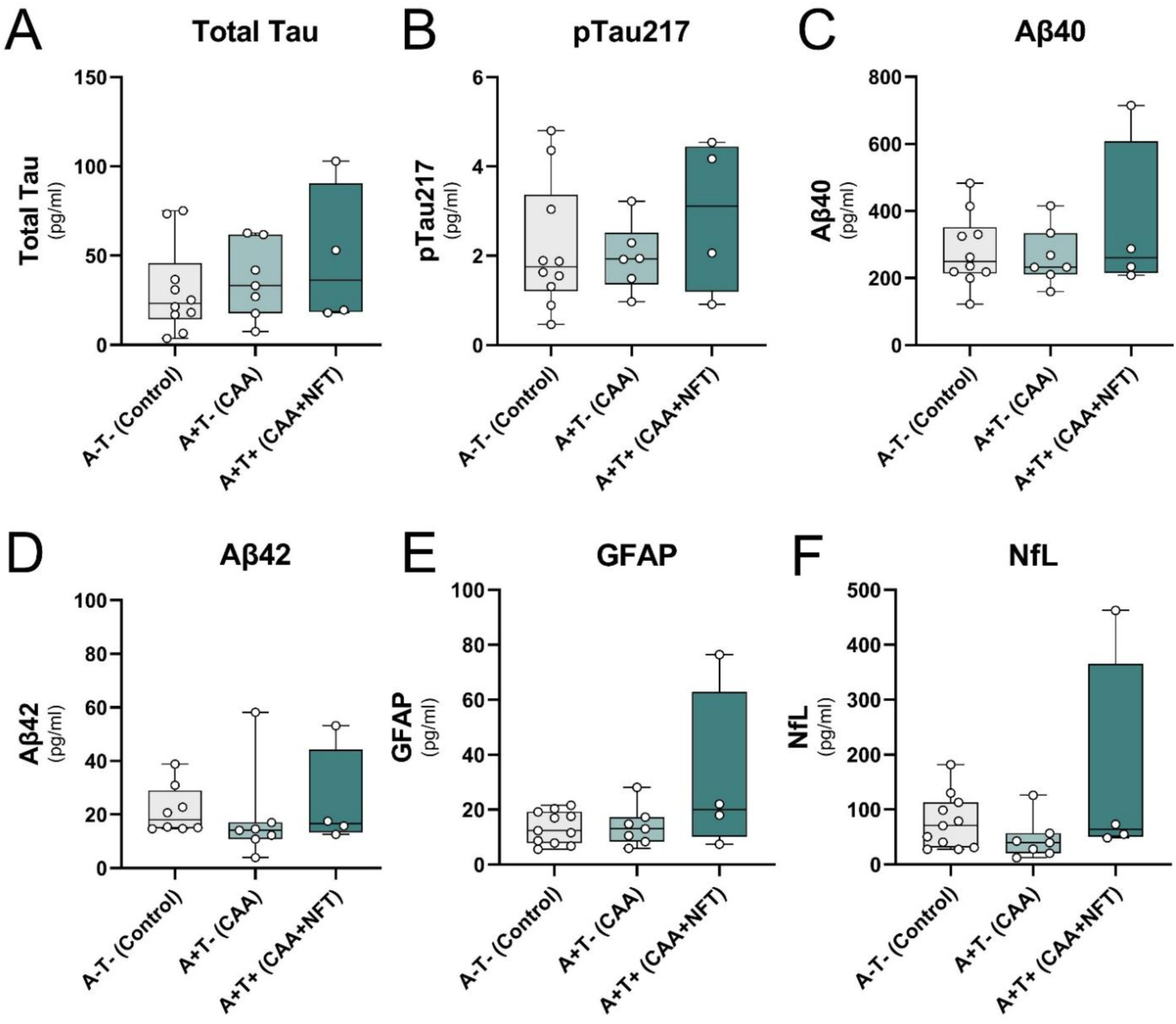
Plasma biomarker levels of total tau (A), phosphorylated tau 217 (pTau217; B), amyloid-beta 40 (Aβ40; C) and 42 (Aβ42; D), glial fibrillary acidic protein (GFAP; E), and neurofilament light (NfL; F) in chimpanzees with (+) and without (-) moderate/severe cerebral amyloid angiopathy (A) and the presence of neocortical neurofibrillary tangles (T). Error bars represent SD.

**Figure 6.**
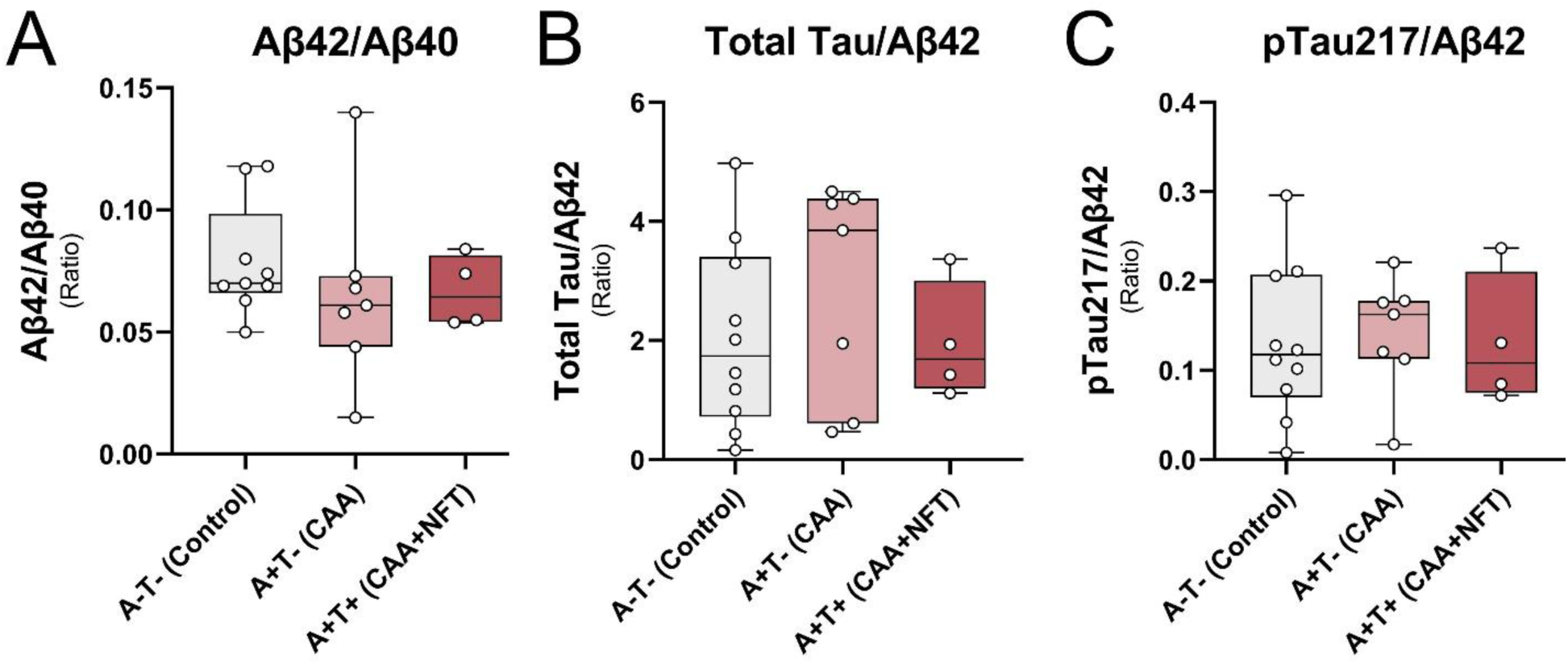
Plasma biomarker ratios of Aβ42/Aβ40 (A), total tau/Aβ42 (B), and pTau217/Aβ42 (C) in chimpanzees with (+) and without (-) moderate/severe cerebral amyloid angiopathy (A) and the presence of neocortical neurofibrillary tangles (T). Error bars represent SD.

**Figure 7.**
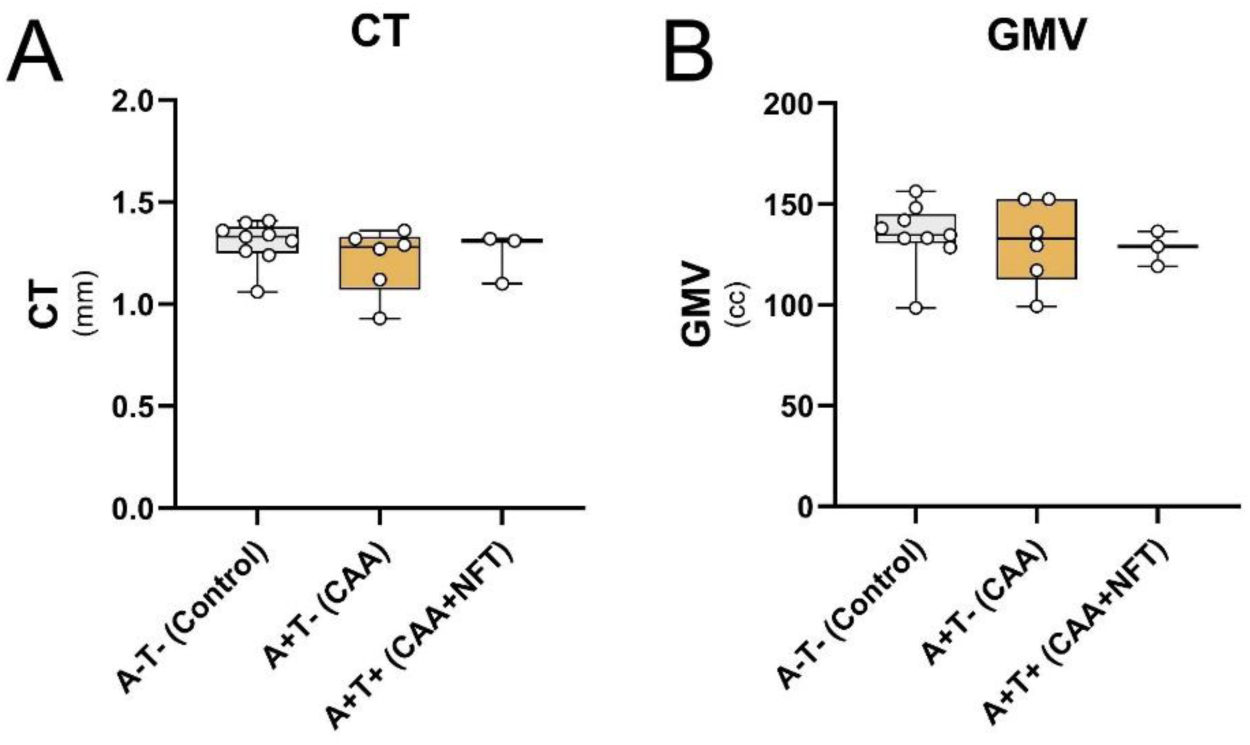
Cortical thickness (CT; A) and gray matter volume (GMV; B) in chimpanzees with (+) and without (-) moderate/severe cerebral amyloid angiopathy (A) and the presence of neocortical neurofibrillary tangles (T). Error bars represent SD.

**Figure 8.**
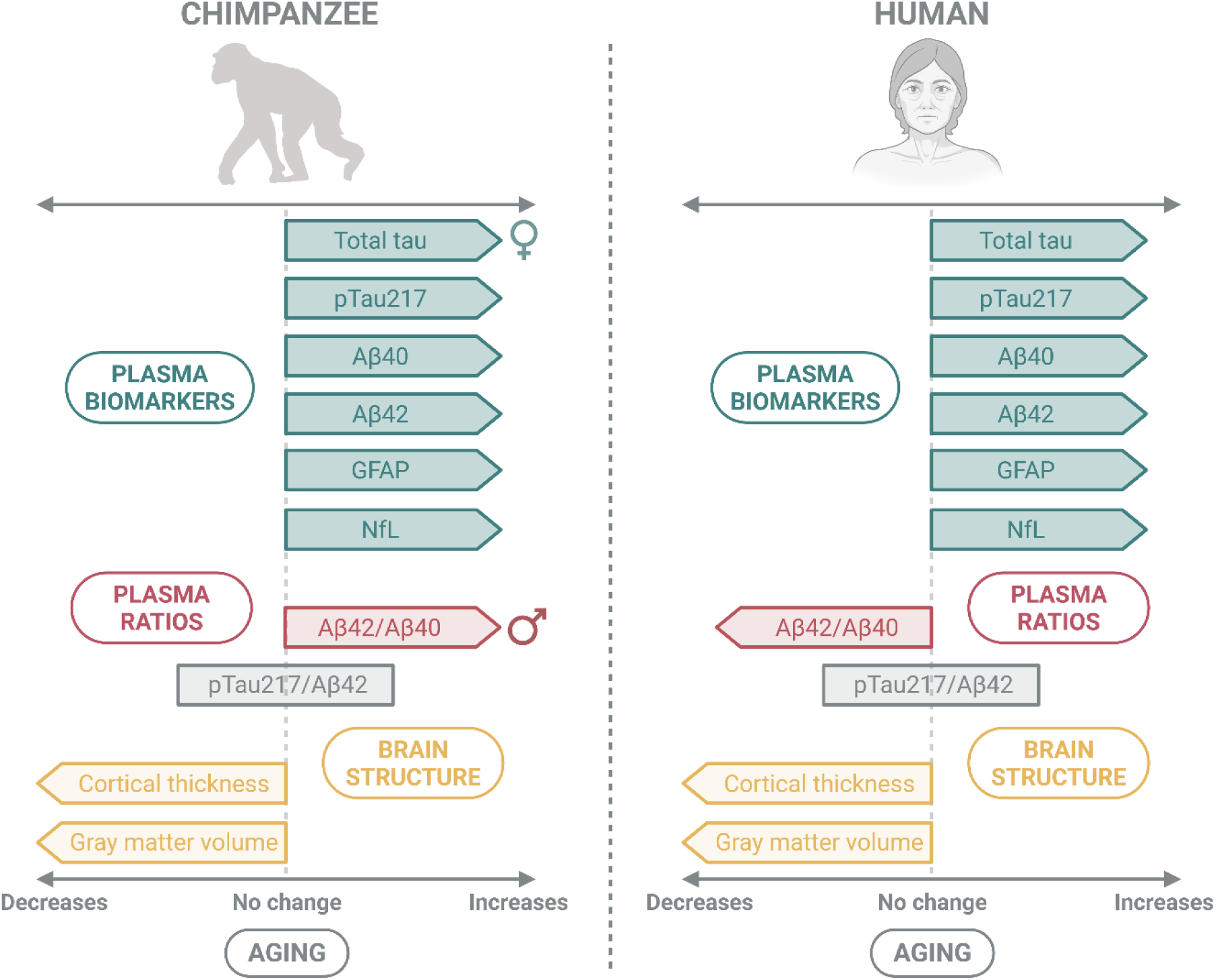
A summary of aging-related changes in AD/ADRD-related plasma biomarkers, plasma ratios, and measures of brain structure in humans and chimpanzees. Created in https://BioRender.com.

We also provide previously reported plasma biomarker reference values in cognitively unimpaired humans or AD individuals collected using the same multiplex assays and Meso Scale Diagnostic platform as was utilized in this investigation to examine possible protein level differences between species (**Table 5**). While these data should be interpreted carefully, a noteworthy difference is higher median/mean levels of plasma GFAP and Aβ40 in humans compared to chimpanzees. GFAP is a protein upregulated during astrogliosis, and it is increased in the plasma of AD patients compared to cognitively intact persons (Benussi et al., 2025; Honey et al., 2022; Santos et al., 2025; Sun et al., 2025; Yu et al., 2025). Intriguingly, our prior work found that chimpanzees do not exhibit an overall increase in GFAP-immunoreactive astrocyte density in association with amyloid plaques and NFT as has been observed in AD brains (Ekonomou et al., 2015; Harpin et al., 1990; Hoozemans et al., 2011; Marlatt et al., 2014; McGeer et al., 1987; Serrano-Pozo et al., 2011; Streit & Xue, 2009). Perhaps, the potentially dampened immune response to AD/ADRD lesions in chimpanzees could be due, in part, to reduced GFAP levels compared to humans. Indeed, elevated plasma GFAP levels in a recent study were associated with increased Aβ and tau burden, reduced cerebral glucose metabolism, and smaller hippocampal volume, as well as poorer cognitive performance in humans (Kim et al., 2026). While species’ variation in plasma GFAP may reflect a distinct contribution to neuroinflammation, higher plasma Aβ40 in humans may signify a greater burden of systemic health conditions, such as cardiovascular and metabolic diseases, potentially influenced by Western diets (Liu et al., 2023; Maurya et al., 2023). For instance, Aβ40 blood concentrations have been associated with cardiac dysfunction, atherosclerosis, metabolic syndrome, and diabetes in humans (Delialis et al., 2024; Janelidze et al., 2016; Sopova et al., 2026; Stamatelopoulos et al., 2018).

**Table 5.**
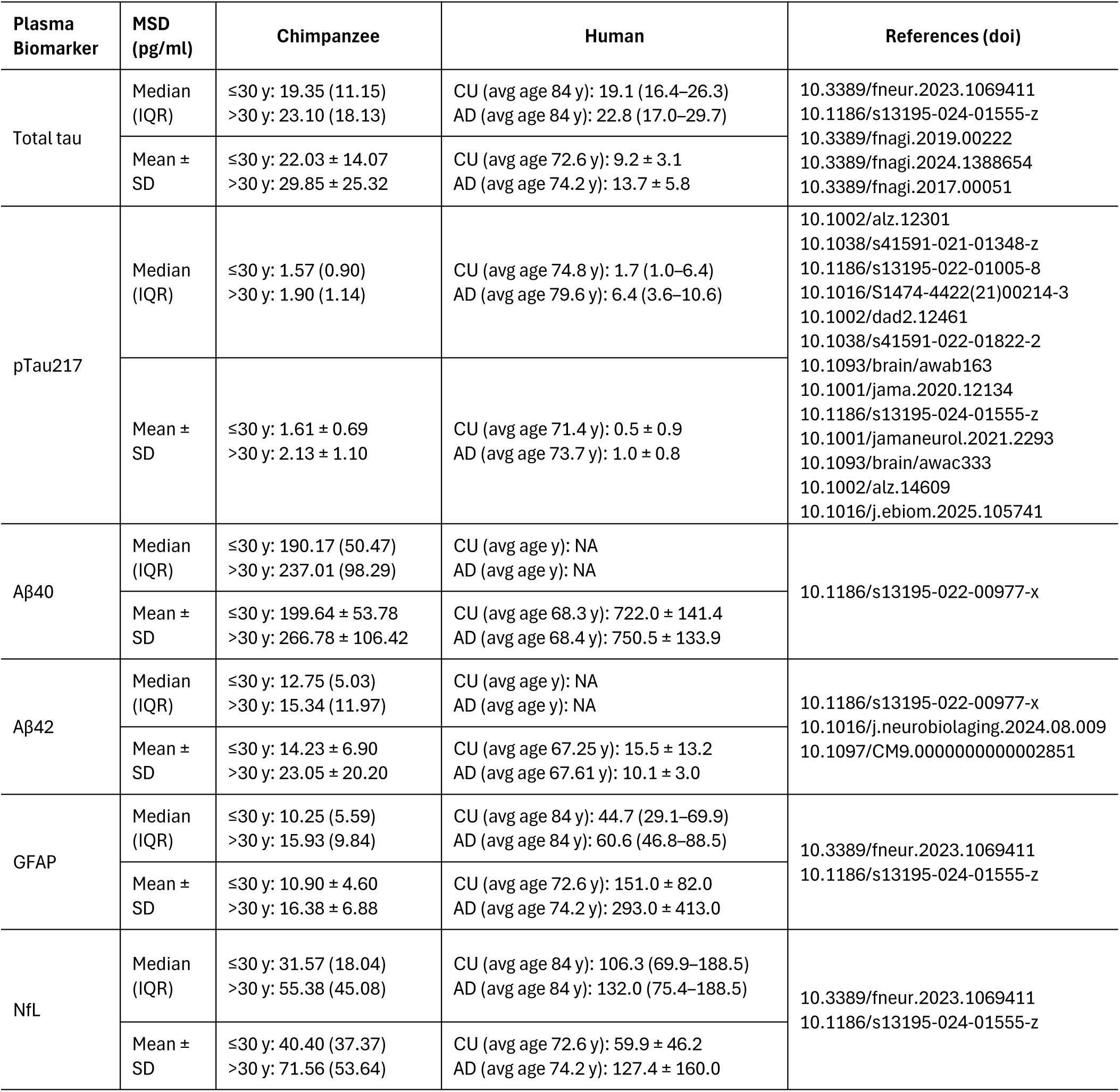
Plasma reference values (pg/ml) by median with interquartile range (IQR) and mean ± standard deviation (SD) in chimpanzees ≤ 30 years (y) or > 30 years of age and humans who are cognitively unimpaired (CU) or have Alzheimer’s disease (AD) using the Meso Scale Diagnostic (MSD) platform.

| **Plasma Biomarker** | **MSD (pg/ml)** | **Chimpanzee** | **Human** | **References (doi)** |
|---|---|---|---|---|
| Total tau | Median (IQR) | ≤30 y: 19.35 (11.15)<br>>30 y: 23.10 (18.13) | CU (avg age 84 y): 19.1 (16.4–26.3)<br>AD (avg age 84 y): 22.8 (17.0–29.7) | 10.3389/fneur.2023.1069411<br>10.1186/s13195-024-01555-z<br>10.3389/fnagi.2019.00222<br>10.3389/fnagi.2024.1388654<br>10.3389/fnagi.2017.00051 |
|  | Mean ± SD | ≤30 y: 22.03 ± 14.07<br>>30 y: 29.85 ± 25.32 | CU (avg age 72.6 y): 9.2 ± 3.1<br>AD (avg age 74.2 y): 13.7 ± 5.8 |  |
| pTau217 | Median (IQR) | ≤30 y: 1.57 (0.90)<br>>30 y: 1.90 (1.14) | CU (avg age 74.8 y): 1.7 (1.0–6.4)<br>AD (avg age 79.6 y): 6.4 (3.6–10.6) | 10.1002/alz.12301<br>10.1038/s41591-021-01348-z<br>10.1186/s13195-022-01005-8<br>10.1016/S1474-4422(21)00214-3<br>10.1002/dad2.12461<br>10.1038/s41591-022-01822-2<br>10.1093/brain/awab163<br>10.1001/jama.2020.12134<br>10.1186/s13195-024-01555-z<br>10.1001/jamaneurol.2021.2293<br>10.1093/brain/awac333<br>10.1002/alz.14609<br>10.1016/j.ebiom.2025.105741 |
|  | Mean ± SD | ≤30 y: 1.61 ± 0.69<br>>30 y: 2.13 ± 1.10 | CU (avg age 71.4 y): 0.5 ± 0.9<br>AD (avg age 73.7 y): 1.0 ± 0.8 |  |
| Aβ40 | Median (IQR) | ≤30 y: 190.17 (50.47)<br>>30 y: 237.01 (98.29) | CU (avg age y): NA<br>AD (avg age y): NA | 10.1186/s13195-022-00977-x |
|  | Mean ± SD | ≤30 y: 199.64 ± 53.78<br>>30 y: 266.78 ± 106.42 | CU (avg age 68.3 y): 722.0 ± 141.4<br>AD (avg age 68.4 y): 750.5 ± 133.9 |  |
| Aβ42 | Median (IQR) | ≤30 y: 12.75 (5.03)<br>>30 y: 15.34 (11.97) | CU (avg age y): NA<br>AD (avg age y): NA | 10.1186/s13195-022-00977-x<br>10.1016/j.neurobiolaging.2024.08.009<br>10.1097/CM9.0000000000002851 |
|  | Mean ± SD | ≤30 y: 14.23 ± 6.90<br>>30 y: 23.05 ± 20.20 | CU (avg age 67.25 y): 15.5 ± 13.2<br>AD (avg age 67.61 y): 10.1 ± 3.0 |  |
| GFAP | Median (IQR) | ≤30 y: 10.25 (5.59)<br>>30 y: 15.93 (9.84) | CU (avg age 84 y): 44.7 (29.1–69.9)<br>AD (avg age 84 y): 60.6 (46.8–88.5) | 10.3389/fneur.2023.1069411<br>10.1186/s13195-024-01555-z |
|  | Mean ± SD | ≤30 y: 10.90 ± 4.60<br>>30 y: 16.38 ± 6.88 | CU (avg age 72.6 y): 151.0 ± 82.0<br>AD (avg age 74.2 y): 293.0 ± 413.0 |  |
| NfL | Median (IQR) | ≤30 y: 31.57 (18.04)<br>>30 y: 55.38 (45.08) | CU (avg age 84 y): 106.3 (69.9–188.5)<br>AD (avg age 84 y): 132.0 (75.4–188.5) | 10.3389/fneur.2023.1069411<br>10.1186/s13195-024-01555-z |
|  | Mean ± SD | ≤30 y: 40.40 (37.37)<br>>30 y: 71.56 (53.64) | CU (avg age 72.6 y): 59.9 ± 46.2<br>AD (avg age 74.2 y): 127.4 ± 160.0 |  |

## Limitations of this study

Although chimpanzees are among the closest living evolutionary relatives of humans, differences in lifespan, aging trajectories, and AD/ADRD-related pathology limit direct translation to human AD/ADRD, and it remains unclear whether the same degree of AD/ADRD-related neuropathology observed in human brains is necessary to induce a comparable level of cognitive impairment in chimpanzees or other NHP (M. K. Edler et al., 2026; Freire-Cobo et al., 2021). Attempts to directly compare clinically defined AD/ADRD-related cognitive impairment in humans with reported cognitive decline in chimpanzees and other NHP is impossible due to the substantial differences in the assessment and psychometric analyses of performance data between species. Furthermore, neurotransmitter changes, alterations in dendritic spine morphology, spine and synapse loss, and myelin alterations occur in humans and NHP, including chimpanzees, macaques, and marmosets, in absence of significant AD-like pathology as a manifestation of normal brain aging (reviewed in Freire-Cobo et al., 2021). Therefore, at this time, plasma biomarker patterns in chimpanzees should be interpreted as comparative profiles of aging and neurodegeneration rather than direct equivalents of human clinical AD/ADRD biomarkers. Secondly, the primary analyses are cross-sectional, which limits our ability to infer within-individual biomarker trajectories or determine whether changes in biomarkers precede cognitive, imaging, or neuropathological outcomes. Additional studies with repeated sampling across longer intervals will be important for determining these temporal changes. Furthermore, some plasma biomarkers are not specific to brain pathology. For example, plasma total tau and pTau217 assays capture circulating tau from both central and peripheral sources and do not specifically measure brain-derived tau. Similarly, inflammatory proteins such as GFAP can be influenced by systemic disease and comorbid conditions. Captive chimpanzees also differ from humans in environment, diet, activity patterns, metabolism, and prevalence of comorbidities such as cardiovascular disease, which may influence circulating biomarkers. Thus, some biomarker differences may reflect systemic physiology, stress, inflammation, or species-specific aging biology rather than progressive AD-like neuropathology alone. Lastly, renal function is an important consideration for interpreting blood-based biomarkers, as impaired clearance can influence circulating protein concentrations. Estimating renal function in chimpanzees is challenging because standard human equations for estimated glomerular filtration rate are not directly validated for this species. Therefore, residual confounding by renal function or species-specific differences in protein clearance cannot be excluded.

## Conclusion

This study is the first to validate and utilize assays to measure biomarkers of aging and AD/ADRD using chimpanzee plasma across a broad age range, although a preliminary report examining serum biomarkers of aging in chimpanzees supports the significant associations between aging, plasma biomarkers, and brain structure (M. Mulholland et al., 2026). In humans, many studies of plasma biomarkers associated with AD/ADRD have focused on midlife and older adults, when pathology is underway. Because aging is the strongest risk factor for AD/ADRD, evaluating biomarker profiles across earlier and later life stages may help identify when neurodegenerative or resilience-associated biological changes first emerge and when interventions may be most effective for preventing or slowing disease progression. Additionally, we validated the multiplex biomarker assays in-house for compatibility with chimpanzee plasma before applying them to the study cohort. This validation step is pertinent as commercially available human biomarker assays may not perform identically in NHP samples, and species-specific differences in protein sequence, matrix effects, or assay interference could affect accuracy.

Chimpanzees provide an invaluable comparative aging model, as they share genetic, physiological, and neurobiological similarity with humans. They also develop extensive Aβ burden, but only express relatively minimal neocortical NFT load, neuroinflammation, and neuronal loss, although whether these neuropathologies translate to decline in cognitive function comparable to humans remains to be fully established in chimpanzees (M. Edler et al., 2023; M. K. Edler et al., 2017, 2018, 2020, 2021, 2026; Freire-Cobo et al., 2021; Munger et al., 2019) . Continuing to investigate aging and AD/ADRD-related changes in humans and NHP is crucial to understanding comparative vulnerability to neurodegeneration and whether AD/ADRD is, or is not, a uniquely human condition. Beyond the translational value of our findings, this research may also inform chimpanzee captive management and care programs. Incorporating a comprehensive fluid biomarker assessment program in the U.S. captive chimpanzee population may benefit their care and welfare by identifying apes at risk or currently experiencing neurodegeneration or other age-related impairments which may impact their quality of life.

## Author Contributions

MKE, IMW, and MAR designed research; MKE and IMW performed research; CCS and JRR contributed samples, reagents, and analytic tools; MKE, IMW, MAR, and JRR analyzed data; MKE, IMW, MAR, CCS, PRH, and JRR wrote the paper.

## Competing Interest Statement

All authors report no biomedical financial interests or potential conflicts of interest.

## Acknowledgements

We thank Dr. Elizabeth Magden for collecting the plasma samples and Dr. William D. Hopkins for generously sharing plasma samples and archival MRI data for this study. This work was supported, in part, by NIH grants AG067419, AG087945, AG073305, NS092988, NS130713, and the Dianne Isakson Distinguished Professorship. Chimpanzee maintenance at the National Center for Chimpanzee Care was previously funded by NIH/NCRR U42-OD-011197. All aspects of this research conformed to existing U.S. and NIH federal policies on the ethical use of chimpanzees in research and was approved by the ORIP Chimpanzee Research User Panel. Additional funds were provided by the Cancer Neuroscience Program at the University of Texas MD Anderson Cancer Center.

## Notes

### Competing Interest Statement

The authors have declared no competing interest.

