## Supplemental Materials for "Plasma biomarkers of Alzheimer’s disease reveal conserved features of age-related neurodegeneration in chimpanzees and humans"

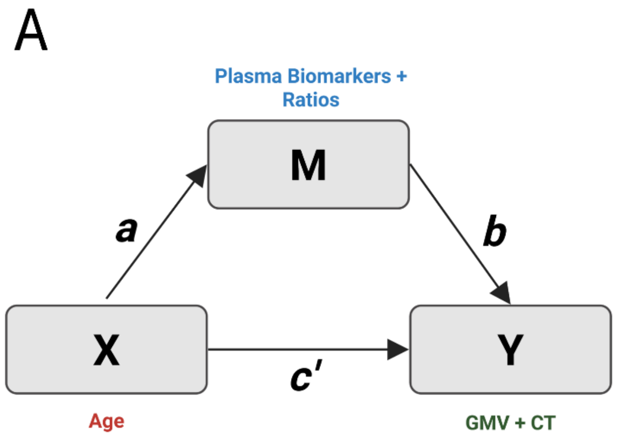


**Figure S1.** Simple mediation utilizes linear regression to measure the total, direct, and indirect effect of the independent variable, X, as well as the mediator, M, on the dependent variable, Y (A, SPSS Hayes Model 4). The correlation between X and M is represented by path "a" and between M and Y by path "b", and the regression coefficient "c" depicts the direct effect of X on Y, whereas coefficients "a" and "b" convey the indirect effect of X on Y. The total effect of X on Y is "c + a*b". Moderated mediation entails regressing the mediator, M, on the independent variable, X, moderator, W, and interaction of X*W for path "a", while path "b" represents regressing the mediator, M, on the dependent variable, Y, and path "c" represents regressing the independent variable, X, on the dependent variable, Y (B, SPSS Hayes Model 7). The moderated mediation model tests for the indirect effect of X on Y via the proposed mediator, M, with the indirect effect being moderated by W. Created in https://BioRender.com.

**Table S1**. Assay analytics and limits of detection for plasma biomarkers in chimpanzees.

| Plasma Biomarkers | Catalog Number | Sample Dilution (Fold) | LOD (pg/ml) | LLOQ (pg/ml) | ULOQ (pg/ml) | Dynamic Range (pg/ml) | Inter-assay precision for internal QC sample (%CV) |
|---|---|---|---|---|---|---|---|
| Aβ40 (pg/ml) | K15200E | 4 | 9.970 | 50.00 | 7,000 | 9.97 - 7,000 | 0.36 |
| Aβ42 (pg/ml) | | | 0.368 | 3.13 | 1,270 | 0.37 - 1,270 | 1.08 |
| GFAP (pg/ml) | K151ANV-2 | 2 | 0.167 | 0.45 | 800 | 0.15 - 880 | 2.38 |
| NfL (pg/ml) | | | 1.210 | 4.53 | 4,330 | 0.93 - 4,330 | 5.77 |
| p-Tau217$_{Lilly}$ (pg/ml) | K151APFS | Neat | 0.695 | 2.10 | 3,590 | 0.88 - 3,590 | 2.56 |
| Total Tau (pg/ml) | K151APSS | Neat | 0.025 | 0.04 | 153 | 0.04 - 153 | 2.85 |

LOD - Limit of Detection; LLOQ - Lower Limit of Quantification; ULOQ - Upper Limit of Quantification

*LOD < LLOQ:* LOD represents the minimum detectable signal, while LLOQ represents the minimum quantifiable level with confidence.

*Dynamic Range:* Spans from LLOQ to ULOQ, where results are accurate and precise.

**Table S2**. Linearity, parallelism, and spike recovery for plasma assays in chimpanzees.

| Plasma Biomarker | Linearity | | | Parallelism | | Spike Recovery | | |
|---|---|---|---|---|---|---|---|---|
| | *Dilutions Tested (Fold)* | *Concentration Range (pg/ml)* | *$R^2$* | *Dilutions Tested (Fold)* | *p* | *Dilutions Tested (Fold)* | *Spike Concentrations (pg/ml)* | *% Recovery* |
| Aβ40 | 0, 2, 3, 4, 6, 8 | 21.21 - 150.89 | 0.991 | 0, 2, 3, 4, 6, 8 | 0.630 | 0, 4, 8 | 0 - Low: 89.92; High: 229.68<br>4 - Low: 27.05; High: 135.81<br>8 - Low: 19.31; High: 120.48 | 0 - Low: 108.25; High: 116.40<br>4 - Low: 88.61; High: 93.80<br>8 - Low: 101.73; High: 90.43 |
| Aβ42 | 0, 2, 3, 4, 6, 8 | 2.69 - 10.38 | 0.997 | 0, 2, 3, 4, 6, 8 | 0.425 | 0, 4, 8 | 0 - Low: 8.42; High: 21.68<br>4 - Low: 2.88; High: 13.02<br>8 - Low: 2.01; High: 12.25 | 0 - Low: 138.75; High: 112.17<br>4 - Low: 93.89; High: 79.78<br>8 - Low: 96.32; High: 79.85 |
| GFAP | 0, 2, 4, 8, 16 | 0.86 - 13.98 | 0.995 | 0, 2, 4, 8, 16 | 0.484 | 4, 8, 16 | 4 - Low: 2.29; High: 7.73<br>8 - Low: 1.26; High: 6.50<br>16 - Low: 0.65; High: 6.71 | 4 - Low: 105.54; High: 88.79<br>8 - Low: 107.83; High: 84.35<br>16 - Low: 75.67; High: 90.72 |
| NfL | 0, 2, 4, 8, 16 | 24.90 - 374.57 | 0.984 | 0, 2, 4, 8, 16 | 0.837 | 4, 8, 16 | 4 - Low: 53.22; High: 77.33<br>8 - Low: 26.51; High: 57.35<br>16 - Low: 14.56; High: 38.63 | 4 - Low: 114.33; High: 97.00<br>8 - Low: 113.89; High: 101.58<br>16 - Low: 80.74; High: 80.74 |
| p-Tau217$_{Lilly}$ | 0, 2, 3, 4, 5, 6 | 0.25 - 1.16 | 0.999 | 0, 2, 3, 4, 5, 6 | 0.358 | 0, 2, 4 | 0 - Low: 2.20; High: 26.21<br>2 - Low: 2.13; High: 25.77<br>4 - Low: 1.88; High: 27.36 | 0 - Low: 93.18; High: 90.52<br>2 - Low: 94.55; High: 89.38<br>4 - Low: 94.89; High: 95.76 |
| Total Tau | 0, 2, 3, 4, 6, 8 | 5.27 - 32.59 | 0.993 | 0, 2, 3, 4, 6, 8 | 0.135 | 0, 2 | 0 - Low: 16.95; High: 18.07<br>2 - Low: 8.91; High: 10.49 | 0 - Low: 103.58; High: 103.34<br>2 - Low: 95.07; High: 100.06 |

*Acceptance Criteria:* Linearity, $R^2 \geq 0.95$ and/or observed concentrations within ±20% of expected; parallelism, lines parallel if $p>0.05$; spike recovery, 80–120% of expected concentration.

**Table S3**. Average plasma biomarker protein levels (pg/ml) by decade (age range: y, years) and sex (M, male, F, female) in chimpanzees. Mean ± standard deviation (SD).

| **Plasma Biomarker** | | **Total Tau** | | **pTau217** | |
|---|---|---|---|---|---|
| *Age Range (y)* | *N (M,F)* | *M* | *F* | *M* | *F* |
| *10-19* | 5, 10 | 27.74 ± 19.75 | 14.27 ± 5.15 | 1.69 ± 0.78 | 1.65 ± 0.60 |
| *20-29* | 32, 38 | 26.29 ± 15.85 | 19.57 ± 12.07 | 1.77 ± 0.76 | 1.43 ± 0.63 |
| *30-39* | 14, 18 | 39.51 ± 36.40 | 32.86 ± 34.48 | 2.28 ± 1.27 | 1.99 ± 1.10 |
| *40-49* | 12, 20 | 25.41 ± 18.99 | 25.60 ± 14.97 | 2.09 ± 1.03 | 1.82 ± 0.89 |
| *>50* | 1, 4 | 59.17 ± 0.00 | 53.33 ± 37.47 | 4.83 ± 0.00 | 3.09 ± 1.07 |
| **Plasma Biomarker** | | **Aβ40** | | **Aβ42** | |
| *Age Range (y)* | *N (M,F)* | *M* | *F* | *M* | *F* |
| *10-19* | 5, 10 | 164.96 ± 11.80 | 207.59 ± 50.86 | 12.23 ± 3.02 | 16.07 ± 5.21 |
| *20-29* | 32, 38 | 203.94 ± 69.48 | 198.46 ± 40.57 | 14.18 ± 6.72 | 14.10 ± 7.78 |
| *30-39* | 14, 18 | 227.96 ± 114.55 | 235.99 ± 94.12 | 13.56 ± 4.93 | 16.69 ± 10.50 |
| *40-49* | 12, 20 | 269.84 ± 74.76 | 293.58 ± 127.03 | 35.95 ± 30.80 | 24.53 ± 19.70 |
| *>50* | 1, 4 | 277.32 ± 0.00 | 292.80 ± 123.94 | 20.88 ± 0.00 | 26.16 ± 19.52 |
| **Plasma Biomarker** | | **GFAP** | | **NfL** | |
| *Age Range (y)* | *N (M,F)* | *M* | *F* | *M* | *F* |
| *10-19* | 5, 10 | 7.45 ± 2.04 | 13.61 ± 4.40 | 41.31 ± 25.52 | 29.38 ± 14.93 |
| *20-29* | 32, 38 | 10.13 ± 4.63 | 11.30 ± 4.53 | 47.84 ± 55.53 | 36.93 ± 18.98 |
| *30-39* | 14, 18 | 12.86 ± 3.99 | 13.30 ± 3.72 | 76.75 ± 67.26 | 60.38 ± 46.11 |
| *40-49* | 12, 20 | 14.91 ± 5.90 | 20.17 ± 8.51 | 68.00 ± 36.30 | 76.51 ± 62.15 |
| *>50* | 1, 4 | 20.31 ± 0.00 | 18.74 ± 7.83 | 167.96 ± 0.00 | 135.09 ± 72.48 |

**Table S4**. Average plasma ratios by decade (age range: y, years) and sex (M, male, F, female) in chimpanzees. Mean ± standard deviation (SD).

| **Plasma Ratio** | | **Aβ42/Aβ40** | | **Total Tau/Aβ42** | |
|---|---|---|---|---|---|
| *Age Range (y)* | *N (M,F)* | *M* | *F* | *M* | *F* |
| *10-19* | 5, 10 | 0.074 ± 0.02 | 0.079 ± 0.01 | 2.269 ± 1.27 | 0.929 ± 0.46 |
| *20-29* | 32, 38 | 0.069 ± 0.02 | 0.070 ± 0.03 | 2.164 ± 1.43 | 1.548 ± 0.92 |
| *30-39* | 14, 18 | 0.070 ± 0.03 | 0.074 ± 0.04 | 2.195 ± 1.79 | 2.133 ± 1.61 |
| *40-49* | 12, 20 | 0.133 ± 0.10 | 0.080 ± 0.04 | 1.020 ± 0.90 | 1.785 ± 1.46 |
| *>50* | 1, 4 | 0.075 ± 0.00 | 0.063 ± 0.02 | 2.834 ± 0.00 | 2.196 ± 0.70 |
| **Plasma Ratio** | | **pTau217/Aβ42** | | | |
| *Age Range (y)* | *N (M,F)* | *M* | *F* | | |
| *10-19* | 5, 10 | 0.144 ± 0.06 | 0.109 ± 0.03 | | |
| *20-29* | 32, 38 | 0.146 ± 0.08 | 0.110 ± 0.06 | | |
| *30-39* | 14, 18 | 0.155 ± 0.10 | 0.144 ± 0.10 | | |
| *40-49* | 12, 20 | 0.083 ± 0.06 | 0.129 ± 0.09 | | |
| *>50* | 1, 4 | 0.231 ± 0.00 | 0.104 ± 0.04 | | |

**Table S5**. Average CT (mm) and whole brain GMV (cc) by decade (age range: y, years) and sex (M, male; F, female) in chimpanzees. Mean ± standard deviation (SD).

| **Cortical Structure** | | **CT** | | **GMV** | |
|---|---|---|---|---|---|
| *Age Range (y)* | *N (M,F)* | *M* | *F* | *M* | *F* |
| *10-19* | 5, 10 | 1.33 ± 0.07 | 1.45 ± 0.04 | 158.59 ± 12.26 | 157.20 ± 9.88 |
| *20-29* | 32, 38 | 1.37 ± 0.08 | 1.39 ± 0.07 | 147.74 ± 12.81 | 150.24 ± 12.74 |
| *30-39* | 14, 18 | 1.33 ± 0.07 | 1.35 ± 0.09 | 147.53 ± 15.11 | 148.05 ± 13.01 |
| *40-49* | 12, 20 | 1.25 ± 0.10 | 1.24 ± 0.15 | 128.98 ± 14.85 | 132.64 ± 21.22 |
| *>50* | 1, 4 | 1.13 ± 0.00 | 1.17 ± 0.17 | 113.32 ± 0.00 | 112.82 ± 19.73 |

**Table S6.** All linear regression statistics for the main effects of age, sex, and interaction of age and sex with plasma biomarkers, plasma ratios, and measures of brain structure

| **MODEL 1: Linear Regression** | | | **Age** | | | **Sex** | | | **Age*Sex Interaction** | | |
|---|---|---|---|---|---|---|---|---|---|---|---|
| *Plasma Biomarkers* | *df* | *Adj $R^2$* | *b* | *t* | *p* | *b* | *t* | *p* | *b* | *t* | *p* |
| Total tau (ln) | 3, 146 | 0.079 | 0.000 | -0.024 | 0.981 | -1.011 | -2.559 | **0.012** | 0.023 | 1.888 | 0.061 |
| pTau217 (ln) | 3, 136 | 0.040 | 0.013 | 1.652 | 0.101 | -0.096 | -0.321 | 0.749 | -0.002 | -0.191 | 0.849 |
| Aβ40 (ln) | 3, 147 | 0.083 | 0.015 | 2.825 | **0.005** | 0.145 | 0.697 | 0.487 | -0.004 | -0.661 | 0.510 |
| Aβ42 (ln) | 3,146 | 0.105 | 0.031 | 3.912 | **<0.001** | 0.494 | 1.617 | 0.108 | -0.018 | -1.917 | 0.057 |
| GFAP (ln) | 3,147 | 0.206 | 0.021 | 3.689 | **<0.001** | 0.272 | 1.212 | 0.227 | -0.004 | -0.502 | 0.617 |
| NfL (ln) | 3, 144 | 0.208 | 0.030 | 3.611 | **<0.001** | -0.193 | -0.590 | 0.556 | 0.002 | 0.160 | 0.873 |
| *Plasma Ratios* | *df* | *Adj $R^2$* | *b* | *t* | *p* | *b* | *t* | *p* | *b* | *t* | *p* |
| Aβ42/Aβ40 (ln) | 3, 145 | 0.036 | 0.016 | 2.706 | **0.008** | 0.465 | 1.969 | **0.051** | -0.018 | -2.438 | **0.016** |
| Total tau/Aβ42 (ln) | 3, 144 | 0.057 | -0.037 | -3.006 | **0.003** | -1.598 | -3.373 | **<0.001** | 0.050 | 3.381 | **<0.001** |
| pTau217/Aβ42 (ln) | 3, 144 | 0.011 | -0.026 | -2.138 | **0.034** | -0.789 | -1.641 | 0.103 | 0.027 | 1.754 | 0.082 |
| *Brain Structure* | *df* | *Adj $R^2$* | *b* | *t* | *p* | *b* | *t* | *p* | *b* | *t* | *p* |
| CT (mm) | 3, 130 | 0.332 | -0.005 | -3.862 | **<0.001** | 0.073 | 1.408 | 0.161 | -0.002 | -0.925 | 0.357 |
| GMV (cc) | 3, 130 | 0.273 | -0.941 | -4.092 | **<0.001** | 2.453 | 0.288 | 0.774 | 0.010 | 0.036 | 0.971 |

| **MODEL 2: Linear Regression** | | | **Age** | | | **Sex** | | |
|---|---|---|---|---|---|---|---|---|
| *Plasma Biomarkers* | *df* | *Adj $R^2$* | *b* | *t* | *p* | *b* | *t* | *p* |
| Total tau (ln) | 2, 148 | 0.063 | 0.016 | 2.655 | **0.009** | -0.299 | -2.517 | **0.013** |
| pTau217 (ln) | 2, 137 | 0.047 | 0.012 | 2.628 | **0.010** | -0.151 | -1.663 | 0.099 |
| Aβ40 (ln) | 2, 148 | 0.086 | 0.012 | 3.968 | **<0.001** | 0.014 | 0.220 | 0.826 |
| Aβ42 (ln) | 2, 147 | 0.089 | 0.018 | 4.065 | **<0.001** | -0.064 | -0.688 | 0.493 |
| GFAP (ln) | 2, 148 | 0.210 | 0.019 | 5.690 | **<0.001** | 0.165 | 2.463 | **0.015** |
| NfL (ln) | 2, 145 | 0.213 | 0.031 | 6.406 | **<0.001** | -0.143 | -1.467 | 0.145 |
| *Plasma Ratios* | *df* | *Adj $R^2$* | *b* | *t* | *p* | *b* | *t* | *p* |
| Aβ42/Aβ40 (ln) | 2, 146 | 0.003 | 0.004 | 1.213 | 0.227 | -0.084 | -1.153 | 0.251 |
| Total tau/Aβ42 (ln) | 2, 145 | -0.011 | -0.003 | -0.394 | 0.694 | -0.069 | -0.473 | 0.637 |
| pTau217/Aβ42 (ln) | 2, 145 | -0.004 | -0.009 | -1.215 | 0.227 | 0.016 | 0.113 | 0.910 |
| *Brain Structure* | *df* | *Adj $R^2$* | *b* | *t* | *p* | *b* | *t* | *p* |
| CT (mm) | 2, 131 | 0.333 | -0.007 | -8.231 | **<0.001** | 0.027 | 1.741 | 0.084 |
| GMV (cc) | 2, 131 | 0.278 | -0.934 | -7.291 | **<0.001** | 2.748 | 1.081 | 0.282 |

Aβ, amyloid-beta, GFAP, glial fibrillary acidic protein, NfL, neurofilament light, pTau217, phosphorylated tau 217

**Table S7.** Plasma biomarker levels (pg/ml) in chimpanzees with (+) and without (-) moderate/severe cerebral amyloid angiopathy (A) and neurofibrillary tangles (T). Mean ± standard deviation (SD).

| **Plasma Biomarker** | | **Total Tau** | **pTau217** | **Aβ40** |
|---|---|---|---|---|
| *Pathology Status* | *N (M,F)* | *Mean ± SD* | *Mean ± SD* | *Mean ± SD* |
| A-T- (Control) | 6, 5 | 30.81 ± 25.00 | 2.18 ± 1.44 | 281.03 ± 107.96 |
| A+T- (CAA) | 2, 5 | 35.99 ± 20.99 | 1.97 ± 0.76 | 264.81 ± 85.39 |
| A+T+ (CAA+NFT) | 0, 4 | 30.22 ± 19.79 | 2.38 ± 1.65 | 243.32 ± 40.27 |
| **Plasma Biomarker** | | **Aβ42** | **GFAP** | **NfL** |
| *Pathology Status* | *N (M,F)* | *Mean ± SD* | *Mean ± SD* | *Mean ± SD* |
| A-T- (Control) | 6, 5 | 21.65 ± 8.97 | 13.53 ± 5.85 | 77.35 ± 49.57 |
| A+T- (CAA) | 2, 5 | 18.72 ± 17.89 | 14.02 ± 7.31 | 46.55 ± 38.01 |
| A+T+ (CAA+NFT) | 0, 4 | 15.31 ± 2.49 | 15.82 ± 7.52 | 58.96 ± 12.56 |

**Table S8.** Plasma biomarker ratios in chimpanzees with (+) and without (-) moderate/severe cerebral amyloid angiopathy (A) and neurofibrillary tangles (T). Mean ± standard deviation (SD).

| **Plasma Ratio** | | **Aβ42/Aβ40** | **Total Tau/Aβ42** | **pTau217/Aβ42** |
|---|---|---|---|---|
| *Pathology Status* | *N (M,F)* | *Mean ± SD* | *Mean ± SD* | *Mean ± SD* |
| A-T- (Control) | 6, 5 | 0.079 ± 0.02 | 2.041 ± 1.56 | 0.131 ± 0.09 |
| A+T- (CAA) | 2, 5 | 0.066 ± 0.04 | 2.866 ± 1.81 | 0.141 ± 0.07 |
| A+T+ (CAA+NFT) | 0, 4 | 0.064 ± 0.02 | 1.970 ± 1.22 | 0.147 ± 0.08 |

**Table S9.** Cortical thickness (CT, mm) and whole brain gray matter volume (GMV, cc) in chimpanzees with (+) and without (-) moderate/severe cerebral amyloid angiopathy (A) and neurofibrillary tangles (T). Mean ± standard deviation (SD).

| **Cortical Structure** | | **CT (mm)** | **GMV (cc)** |
|---|---|---|---|
| *Pathology Status* | *N (M,F)* | *Mean ± SD* | *Mean ± SD* |
| A-T- (Control) | 6, 5 | 1.30 ± 0.11 | 134.79 ± 16.07 |
| A+T- (CAA) | 2, 5 | 1.22 ± 0.16 | 131.24 ± 20.71 |
| A+T+ (CAA+NFT) | 0, 4 | 1.24 ± 0.12 | 128.34 ± 8.69 |

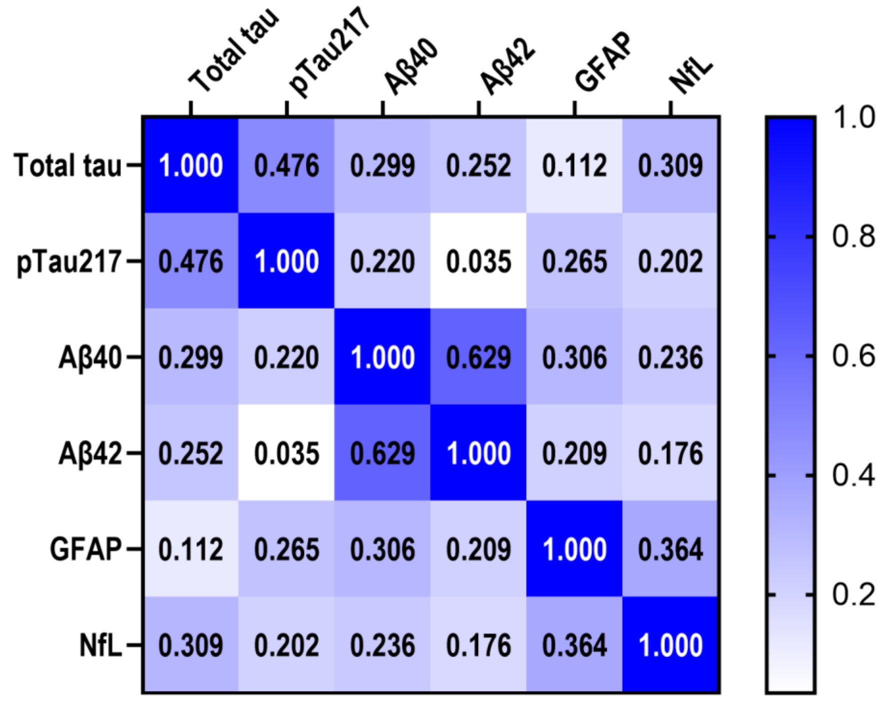


**Figure S2**. Pearson's r correlation coefficients between plasma biomarkers while controlling for age and sex

**Table S10.** P-values for partial correlational analyses of covariation between plasma biomarkers while controlling for age and sex

| **Plasma Biomarker** | *Total Tau* | *pTau217* | *Aβ40* | *Aβ42* | *GFAP* | *NfL* |
|---|---|---|---|---|---|---|
| Total Tau | - | **<0.001** | **<0.001** | **0.004** | 0.212 | **<0.001** |
| pTau217 | **<0.001** | - | **0.013** | 0.699 | **0.003** | **0.022** |
| Aβ40 | **<0.001** | **0.013** | - | **<0.001** | **<0.001** | **0.008** |
| Aβ42 | **0.004** | 0.699 | **<0.001** | - | **0.018** | **0.047** |
| GFAP | 0.212 | **0.003** | **<0.001** | **0.018** | - | **<0.001** |
| NfL | **<0.001** | **0.022** | **0.008** | **0.047** | **<0.001** | - |
